# senSCOPE: Modeling radiative transfer and biochemical processes in mixed canopies combining green and senescent leaves with SCOPE

**DOI:** 10.1101/2020.02.05.935064

**Authors:** Javier Pacheco-Labrador, Tarek S. El-Madany, Christiaan van der Tol, M. Pilar Martin, Rosario Gonzalez-Cascon, Oscar Perez-Priego, Jinhong Guan, Gerardo Moreno, Arnaud Carrara, Markus Reichstein, Mirco Migliavacca

**Affiliations:** Max Planck Institute for Biogeochemistry, Hans Knöll Straße 10, Jena, D-07745, Germany; Faculty of Geo-Information Science and Earth Observation (ITC), University of Twente, PO Box 217, AE Enschede 7500, The Netherlands; Environmental Remote Sensing and Spectroscopy Laboratory (SpecLab), Institute of Economic, Geography and Demography (IEGD-CCHS), Spanish National Research Council (CSIC), C/Albasanz 26-28, 28037 Madrid, Spain; Department of Environment, National Institute for Agriculture and Food Research and Technology (INIA), Ctra. Coruña, Km. 7,5, 28040 Madrid, Spain; Department of Biological Sciences Macquarie University, 6 Wally’s Walk, NSW 2109, Australia; State Key Laboratory of Soil Erosion and Dryland Farming on the Loess Plateau, Northwest A&F University, Yangling, Shaanxi 712100, China; Forest Research Group - INDEHESA University of Extremadura, 10600 Plasencia, Spain; Fundación Centro de Estudios Ambientales del Mediterráneo (CEAM), Charles Darwin 14, Parc Tecnològic, 46980 Paterna, Spain

## Abstract

Semi-arid grasslands and other ecosystems combine green and senescent leaves featuring different biochemical and optical properties, as well as functional traits. Knowing how these properties vary is necessary to understand the functioning of these ecosystems. However, differences between green and senescent leaves are not considered in recent models representing radiative transfer, heat, water and CO_2_ exchange such as the Soil-Canopy Observation of Photosynthesis and Energy fluxes (SCOPE). Neglecting the contribution of senescent leaves to the optical and thermal signal of vegetation limits the possibilities to use remote sensing information for studying these ecosystems; as well as neglecting their lack of photosynthetic activity increases uncertainty in the representation of ecosystem fluxes. In this manuscript we present senSCOPE as a step towards a more realistic representation of mixed green and senescent canopies. senSCOPE is a modified version of SCOPE model that describes a canopy combining green and senescent leaves with different properties and function. The model relies on the same numerical solutions than SCOPE, but exploits the linear nature of the scattering coefficients to combine optical properties of both types of leaf. Photosynthesis and transpiration only take place in green leaves; and different green and senescent leaf temperatures are used to close the energy balance. Radiative transfer of sun-induced fluorescence (SIF) and absorptance changes induced by the xanthophyll cycle action are also simulated. senSCOPE is evaluated against SCOPE both using synthetic simulations, forward simulations based on observations in a Mediterranean tree-grass ecosystem, and inverting dataset of ground measurements of reflectance factors, SIF, thermal radiance and gross primary production on a heterogeneous and partly senescent Mediterranean grassland. Results show that senSCOPE outputs vary quite linearly with the fraction of green leaf area, whereas SCOPE does not respond linearly to the effective leaf properties, calculated as the weighted average of green and senescent leaf parameters. Inversion results and pattern-oriented model evaluation show that senSCOPE improves the estimation of some parameters, especially chlorophyll content, with respect SCOPE retrievals during the dry season. Nonetheless, inaccurate knowledge of the optical properties of senescent matter still complicates model inversion. senSCOPE brings new opportunities for the monitoring of canopies mixing green and senescent leaves, and for improving the characterization of the optical properties of senescent material.

## 1 Introduction

Consistent monitoring of relevant vegetation properties is an essential step towards understanding the response of vegetation function (e.g., photosynthesis, transpiration) to changes in environment. Among others, photosynthetic performance and water use efficiencies are key elements to predict and understand vegetation responses to the climate change scenarios (e.g., elevated atmospheric CO_2_ concentration, higher temperatures and altered water regimes) (IPCC 2014). However, current Land Surface Models (LSM) predictions of these fluxes include large uncertainties (Friedlingstein et al. 2014); partly due to inadequate representation of different processes as well as to the lack of knowledge of functional parameters describing plant function (e.g., maximum carboxylation rate (*V*_cmax_), maximum electron transport rate (*J*_max_) or the Ball-Berry stomatal sensitivity (*m*)) (Rogers 2014; Rogers et al. 2016; Schaefer et al. 2012). Recent efforts of the Remote Sensing (RS) community have focused on the estimation of these parameters either using statistical approaches (Serbin et al. 2015; Silva-Perez et al. 2018), or combining Radiative Transfer Models (RTM) with Soil-Vegetation-Atmosphere Models (SVAT) (Bayat et al. 2018; Camino et al. 2019; Dutta et al. 2019; Pacheco-Labrador et al. 2019; Zhang et al. 2014; Zhang et al. 2018), notably using the Soil-Canopy Observation Photosynthesis and Energy fluxes (SCOPE) model (van der Tol et al. 2009).

SCOPE represents radiative transfer of optical and thermal infrared radiation (TIR), in a homogeneous 1-D canopy, which is coupled with an energy balance and a photosynthesis models predicting heat and water fluxes and carbon assimilation. SCOPE also propagates leaf level sun-induced chlorophyll fluorescence (SIF) emission and absorptance changes related with the activation of the xanthophyll cycle (Vilfan et al. 2018) to top of the canopy radiances. SCOPE uses Fluspect to model leaf optical properties (Vilfan et al. 2016; Vilfan et al. 2018) and combines 4 different canopy RTM representing outgoing radiation (RTM_o_), SIF (RFM_f_, (van der Tol et al. 2016)), TIR emission (RTM_t_) and xanthophyll absorption (RTM_z_, (Vilfan et al. 2018)) that rely on the four stream SAIL extinction and scattering coefficients model (Verhoef 1984). In addition, Yang and colleagues (2017) developed mSCOPE, an extension of SCOPE that uses a different numerical solution of the radiative transfer problem to represent 1-D but vertically heterogeneous canopies.

A current limitation of SCOPE is the lack of representation of within-canopy heterogeneity of vegetation properties, and specifically the separation of green and senescent leaves, which feature large differences in biophysical properties and function. When leaves senesce, flavonoids undergo enzymatic oxidation processes within the leaf producing diverse semiquinones and quinones which can suffer non-enzymatic secondary reactions with phenols, amino acids and proteins or other polyphenols (Pourcel et al. 2007; Taranto et al. 2017). The result is a heterogeneous mixture of complex brown polymers, difficult to characterize *in vivo* and responsible of the yellow and brown tones that these leaves exhibit (Guyot et al. 1996; Pourcel et al. 2007). The characterization of the optical properties of these “senescent” or “brown” pigments of leaves were addressed by Jacquemoud (1988) using albino corn leaves; however, the authors stated that the characterization had to be improved. In fact, the absorption coefficients currently used by Prospect are usually attributed to F. Baret, via personal communication (e.g., (Féret 2009)). Thus, the characterization of senescent pigments is not as thoroughly documented as for other pigments (Feret et al. 2008; Féret et al. 2017; Jacquemoud and Baret 1990; Vilfan et al. 2018), and their concentration is presented in arbitrary units due to the measurement technique used in their determination (Jacquemoud 1988). Also, when leaves further degrade their color changes (Kidnie et al. 2015), and some of their optical properties might vary with respect to those characterized and used by leaf-level RTM. For example, Melendo-Vega et al, (2018) suggested that overestimation of near infrared reflectance factors in a semi-arid grassland could be related to senescent material, and that this effect increased with its longevity.

Commonly used models such as PROSAIL (Jacquemoud et al. 2009), or more recently SCOPE, allocate all the pigments in a unique “effective” according to the averaged concentrations of the different leaves of the canopy. However, this approach does not adequately represent mixed canopies with varying fractions of green and senescent leaves. The presence of non-photosynthetic elements in the canopy has been already addressed in turbid medium RTM (Bach et al. 2001; Braswell et al. 1996; Wenhan 1993) and used to improve the estimation of biophysical parameters such as leaf area index (*LAI*) or chlorophyll concentration (*C*_ab_) or the fraction of absorbed photosynthetically active radiation (Houborg and Anderson 2009; Houborg et al. 2009; Houborg and McCabe 2016; Wenhan 1993). However, senescent and green leaves do not only feature different optical properties, but also different physiological processes. For example, senescent leaves present little or no chlorophyll content (Hörtensteiner 2006; Whitfield and Rowan 1974) so that they do not assimilate CO_2_ through photosynthesis. Also, senescent leaves do not transpire water and lack of stomatal regulation. Senescent leaves can pose problems for the retrieval of biophysical variables if not adequately represented (Bacour et al. 2002; Houborg and Boegh 2008; Wang et al. 2005). Analogously, inadequate representation of green and senescent leaf pools could also potentially induce uncertainties in the simulation of processes at canopy scale related to photosynthesis and transpiration. Finally, given that SCOPE is now widely used for retrieval of functional properties (Bayat et al. 2018; Camino et al. 2019; Dutta et al. 2019; Pacheco-Labrador et al. 2019; Zhang et al. 2014; Zhang et al. 2018), these uncertainties can propagate in the estimated parameters (Pacheco-Labrador et al. 2019). This fact may limit the application of recent approaches combining RTM and SVAT models for the study of canopies or ecosystems featuring large fractions of dry leaves (in particular in grasslands or semi-arid ecosystems) or for the monitoring of vegetation health, crop productivity and phenology.

Senescent material is present in all vegetation, and for a remote sensing perspective is very critical for annual plants such as grasslands (Houborg et al. 2009; Melendo-Vega et al. 2018), which cover about 40% of the Earth’s terrestrial surface (Anderson 2006). Grassland’s phenology and function are strongly governed by water availability, temperature, herbivory, fire, nitrogen deposition or CO_2_ concentration increase (Anderson 2006; Cleland et al. 2006; Figueroa and Davy 1991; Luo et al. 2018; Migliavacca et al. 2011; Richardson et al. 2013). Green plants transit to senescent, recently dead, and long-term dead plants, each of them featuring different biophysical and optical properties (Kidnie et al. 2015). This transition varies with meteorology (Ren and Zhang 2018), biophysical properties (Henry et al. 2008; Sanaullah et al. 2010; Yuan and Chen 2009), plant functional types (Henry et al. 2008) and changes for different parts of the plant (Henry et al. 2008; Koukoura et al. 2003). Usually, even in grasses, leaves fall while stems stay longer and degrade more slowly due to differences in biochemical composition. Therefore, in multi-species grasslands senescence and degradation can take place at different rates and periods, increasing the variability of surface biophysical and optical properties as well as the complexity of modeling and characterization. In fact, senescent material and litter are nowadays considered a challenge for the estimation of biophysical properties in semi-arid grasslands (He and Mui 2010).

In this work, we present senSCOPE, a modified version of the SCOPE model that separates RTM and physiological processes (photosynthesis and transpiration) for green and senescent leaves. senSCOPE aims at improving the representation of radiative transfer and physiology in senescent canopies. The model is then evaluated in three ways:

1. We run a sensitivity analysis comparing forward simulations of SCOPE and senSCOPE under different meteorological conditions and under different combinations of vegetation parameters for different abundances of senescent leaves.
2. We use observations of model parameters and meteorological data at ecosystem scale to predict fluxes and compare them with EC data.
3. We invert SCOPE and senSCOPE against the same dataset of ground observations of carbon fluxes, reflectance factors (*R*), SIF and TIR radiance used by Pacheco-Labrador et al, (2019) for comparison.

As in the former work, functional and biophysical model parameters are estimated by inverting SCOPE against different combinations of the abovementioned variables sampled at plot scale in a Mediterranean grassland. New inversion boundaries are used according to observations of some of the parameters in the site. Results of both inversions are compared and evaluated using pattern-oriented model evaluation approach (Pacheco-Labrador et al. 2019).

## 2. Description of senSCOPE

The model senSCOPE extends the 1-D model SCOPE to describe homogeneous canopies combining green and senescent leaves randomly mixed. Fig. 1 summarizes the conceptual differences between SCOPE and senSCOPE. Green leaves contain chlorophylls and other photosynthetic pigments that allow them to photosynthesize; and regulate their temperature via transpiration. In contrast, senescent leaves only contain senescent pigments and neither photosynthesize nor transpire. These leaves present some microbial activity related to its degradation, and superficial water (e.g., intercepted rainfall) can evaporate from their surface; however these processes are not represented neither by SCOPE nor senSCOPE. In senSCOPE, the leaf RTM Fluspect (Vilfan et al. 2016; Vilfan et al. 2018) simulates reflectance, transmittance and, in the case of green leaves, also fluorescence according to the biochemical and structural properties of each leaf type. Canopy RTM_o_ implemented in SCOPE (van der Tol et al. 2009) is modified to separately compute the radiation absorbed by each leaf type; and the energy balance model is customized to account for the presence of leaves that neither photosynthesize nor transpire. Since green and senescent leaves feature different radiative balances, a modified RTM_t_ model quantifies thermal emission of each of these two leaf types separately and combines them according to the corresponding fractions of leaf area (*f*); then the model calculates scattering and absorption of this diffuse flux. Eventually, fluorescence emission and optical changes induced by the activation of the xanthophyll cycle in the green leaves is propagated to top of canopy (TOC) radiances and *R* using the RTM_f_ (van der Tol et al. 2016) and RTM_z_ (Vilfan et al. 2018) models already implemented in SCOPE. Both models are coded in Matlab (Matwoks Inc., Natick, MA, USA).

**Figure 1.**
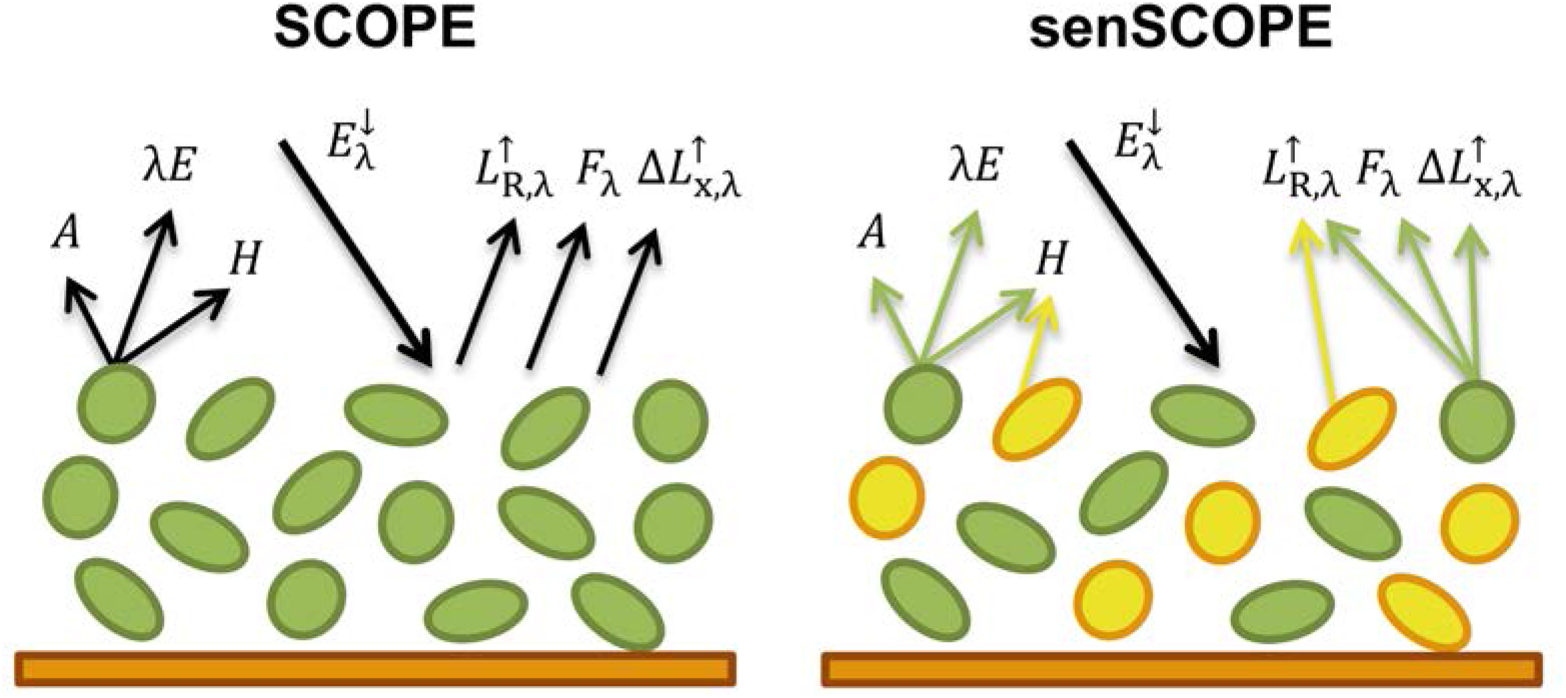
Conceptual differences between SCOPE and senSCOPE models. Green and yellow colours correspond to green and senescent leaves, respectively. Black arrows show processes featured by all leaves; whereas coloured arrows refer to processes featured only for a given type of leaf. The scheme represents assimilation (*A*), latent (*λE*) and sensible heat (*H*) fluxes, incoming spectral irradiance (*E*_*λ*_), reflected spectral radiance (*L*_R,*λ*_), emitted fluorescence radiance (*F*_*λ*_) and changes in *L*_R,*λ*_ due to activation of xanthophyll cycle (*ΔL*_x,*λ*_)

senSCOPE relies on the same solution of the radiative transfer problem implemented in SCOPE, since it exploits the linear nature of the single leaf scattering efficiency factors (Verhoef 1984) to combine the optical properties of green and senescent leaves in an “averaged” leaf. This is simple if leaf angle distribution is assumed the same for both types of leaves. The main advantage of this approach is that it allows representing physiological processes separately in each leaf type. This is important since photosynthesis and transpiration are non-linearly related with radiation and leaf temperature, and therefore might not be adequately represented by a model featuring a unique leaf type characterized by the “averaged parameters” of photosynthetic and non-photosynthetic leaves.

senSCOPE requires a larger number of parameters than SCOPE, since two different leaves must be described, as well as their respective area fractions. Alternatives to minimize the number of parameters and simplify the application of the model in inverse problems are presented in Sect. 3.2.2 and discussed later.

### 2.1 Radiation Fluxes

As SCOPE, senSCOPE relies on SAIL 4-stream theory that can be summarized by a system of four linear equations describing the radiative transfer of the solar direct flux (*E*_s_), the downward diffuse flux (*E*^-^), the upward diffuse flux (*E*^+^), and the flux in the observation direction (*E*_o_).

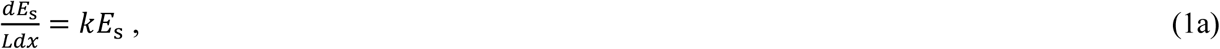

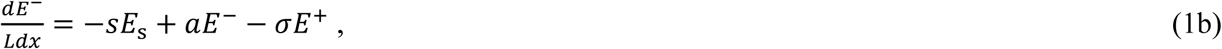

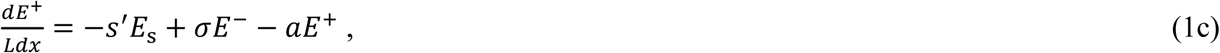

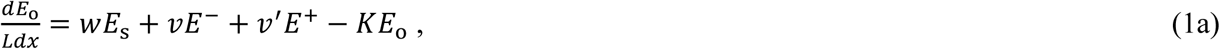

In this system, *x* represents the vertical relative height within the canopy (*x* = 0 for top, *x* = -1 for bottom), and *L* represents the Leaf Area Index (also *LAI*). The remaining variables are the SAIL coefficients defined for first time by Verhoef (1984). *k* and *K* are the extinction coefficients in the solar and observation directions, respectively. They depend on the sun-view geometry, *LAI* and the leaf angle distribution (*LAD*); and they are therefore independent of leaf optical properties. *s, a, σ, s′, w, v* and *v′* are the scattering coefficients depending on sun-view geometry, canopy structure and leaf optical properties. These coefficients define the relationship between a given incident flux (*E*_1_) and a given scattered flux (*E*_2_) in the canopy, and they are computed by integrating single-leaf scattering efficiency factors (*Q*_sc_) that represent the analogous relationship for individual leaves. The scattering coefficient (*b*) corresponding to all the leaves of given zenith inclination angle (*θ*_l_) can be defined as (Verhoef 1984):

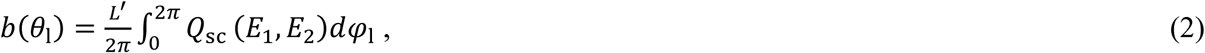

where *L*′ is the *LAI* contained in a horizontal layer of the canopy of width *dx* and *φ*_l_ is the leaf azimuth angle.

As in SCOPE, senSCOPE solves the radiative transfer problem numerically, defining a discrete number of canopy layers and leaf angles. *Q*_sc_(*E*_1_, *E*_2_) are defined assuming that individual leaves are Lambertian diffusors of known hemispherical reflectance (*ρ*) as and hemispherical transmittance (*τ*). *ρ* and *τ* are predicted in SCOPE by Fluspect (Vilfan et al. 2016). For each pair of incident and scattered fluxes, *Q*_sc_(*E*_1_, *E*_2_) is defined as a linear combination of *ρ* and/or *τ* weighted by spectrally invariant factors determined by the geometry of the leaf, or more specifically, the projection of the leaves with respect to the incident flux (*E*_1_) and the downward (-) or upward (+) scattered flux (*E*_2_). As proposed by Bach et al, (2001), senSCOPE exploits this linear nature of *Q*_sc_ to combine the *ρ* and the *τ* of green and senescent leaves into an averaged factors; weighted by their corresponding fractions of leaf area (Eq. 3 and 4). This approach allows applying the solution already proposed by van der Tol et al., (2009) for the linear system shown in Eq. 1a-d.

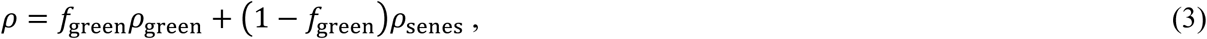

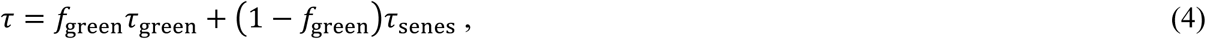

where subscripts “green” and “senes” indicate the type of leaf. Notice that the weighted average of *ρ* and *τ* is not equivalent to the factors predicted for a weighted average of the leaf parameters.

This approach is suitable to represent the radiative transfer of a canopy of homogeneously mixed green and senescent leaves. In order to represent physiological processes for each leaf type separately, it is necessary differentiating the amount total radiation absorbed by each leaf type, and the photosynthetically active radiation (*PAR*) absorbed by chlorophyll (*E*_ap,Chl_). SCOPE quantifies *E*_ap,Cab_ (W m^-2^) using the relative absorption of this pigment respect to the remaining total absorption in the leaf in each spectral band (*k*_Chl,rel_). *E*_ap,Chl_ is computed for the direct (*E*_ap,Chl,dir_) and the diffuse irradiances (*E*_ap,Chl,dif_) as follows:

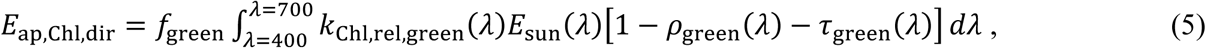

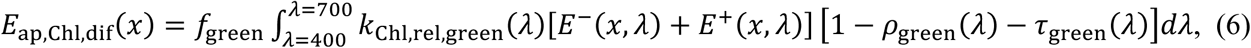

where *λ* is the wavelength and *k*_Chl,rel,green_ is *k*_Chl,rel_ in the green leaves. These quantities are calculated from the upward and downward fluxes without modifying the transfer of radiation. Since senSCOPE defines senescent leaves as containing no chlorophyll, *k*_Chl,rel_ = 0 in senescent leaves and for this reason, absorbed PAR used to simulate photosynthesis in sunlit (*E*_ap,Chl,u_) and shaded leaves (*E*_ap,Chl,h_) per total leaf area of the mixed canopy scales with *f*_green_. Shaded leaves (subscript ‘h’) are only illuminated by diffuse light (Eq. 7); whereas Eq. 5 and 6 must be combined to get *E*_ap,Chl_ in the sunlit leaves (*E*_ap,Chl,u_, subscript ‘u’) (Eq. 8).

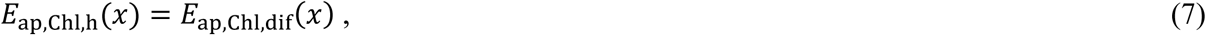

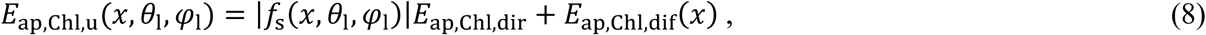

where *f*_s_ is a geometric factor accounting for the projection of each leaf towards the sun.

Total absorbed radiation is used to compute the radiation budget in the canopy and determines leaf temperature, which has implications for photosynthesis and transpiration, and must therefore be computed separately for each leaf type. Total absorbed radiation is computed by SCOPE similarly as in Eq. 5-8, but integrating the fluxes in the full spectral domain (e.g., 400-50.000 nm):

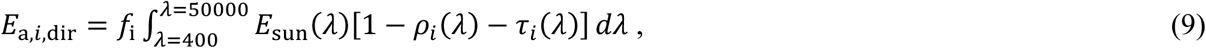

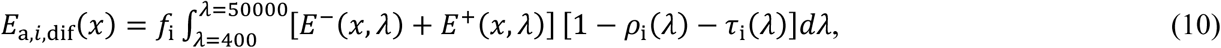

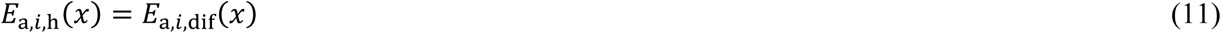

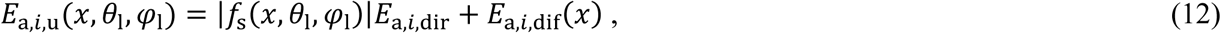

Where subscript “*i*” now stands for either ‘green’ or ‘senescent’.

### 2.2 Energy balance

As in SCOPE, energy balance is closed iteratively by modifying canopy and soil temperatures until the following is met for the soil and for all leaf angles and layers separately:

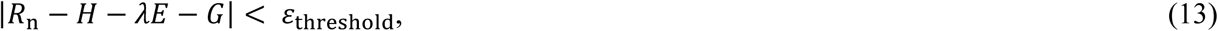

where *R*_n_ is net radiation, *H* is latent heat flux, λ*E* is energy heat flux, *G* is soil heat flux and *ε*_treshold_ is a predefined threshold for the accepted energy balance closure error (*ε*_treshold_), all in W m^-2^.

senSCOPE addresses the energy balance separating the processes occurring in green and senescent leaves, where only the first are assumed to photosynthesize and transpire. Therefore, *ε*_ebal_ is separated into the following elements (Eq. 14):

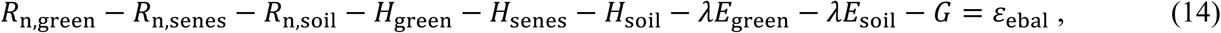

where the subscript “soil” refers to soil fluxes, and only green leaves and soil contribute to λ*E*. However, notice that similarly as in SCOPE, the energy balance is separately closed for soil and for all leaf angles, layers and leaf types.

In order to compute *R*_n_, the contribution of thermal emission must be added to the absorbed radiation calculated in Sect. 2.1. senSCOPE separately represents the temperatures of green and senescent leaves (*T*_c,green_, *T*_c,senes_, respectively) since they absorb radiation cool down differently. Distinguishing these temperatures has an impact on the calculation of photosynthesis, which is temperature dependent. Consequently, black-body thermal emission (*H*_c_) is different for each leaf type (*H*_c,green_, *H*_c,senes_); and the on-sided black-body thermal emission of all leaves is computed as a linear combination of the emission of each leaf type in the canopy:

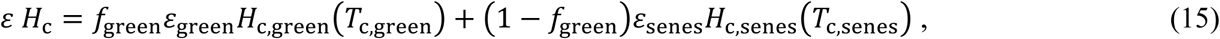

where *ε* is the emissivity, and equals absorptance (1-*ρ*-*τ*) according to Kirchhoff’s Law. The propagation of emitted radiation by leaves and soil through the canopy is calculated using the averaged layer properties as in the original SCOPE. In order to quantify the net thermal radiation (emitted minus absorbed) (*R*_*n*,t_) senSCOPE calculates the amount of energy absorbed by each leaf type using their respective emissivity:

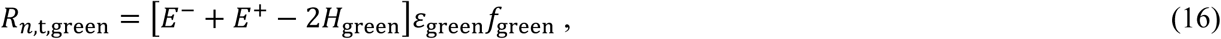

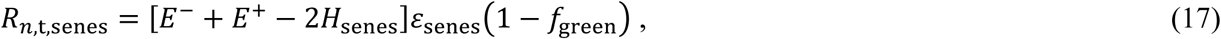

where and *E*^-^ and *E*^+^ are the diffuse emitted fluxes. *R*_*n*,t_ of sunlit and shaded leaves is computed separately. These are energy fluxes per total (senescent plus green) leaf surface area. Therefore, canopy net radiation is computed as the addition of *E*_a_ and *R*_*n*,t_; and *R*_*n*,t_ = *R*_*n*,t,green_ + *R*_*n*,t,senes_ without the need to further weight by fraction.

Aerodynamic resistances are computed as in SCOPE for the whole mixed canopy, since they depend on meteorology and canopy structure. Consequently water and heat fluxes (*H*_green_, *H*_senes_ and λ*E*_green_) in senSCOPE are computed with an identical representation of resistances as in SCOPE, but with leaf temperatures differentiated per leaf type. These fluxes are defined per unit of leaf-type surface, and need to be scaled to the fraction of *LAI* represented by each leaf type in the mixed canopy. Eventually, senSCOPE iteratively resolves six temperatures: sunlit and shaded green leaves (*T*_c,u,green_, *T*_c,h,senes_), sunlit and shaded senescent leaves (*T*_c,u,senes_, *T*_c,h,usenes_), and both sunlit and shaded soil (*T*_s,u_, *T*_s,h_).

### 2.3 Photosynthesis

In senSCOPE, only green leaves photosynthesize and transpire. Photosynthesis is driven by the *PAR* absorbed by chlorophyll (*APAR*_Chl_; which equals *E*_ap,Chl_ transformed from W m^-2^ to μmol m^-2^ s^-1^). The absorbed *PAR* by chlorophyll in green leaves per unit green leaf area is *E*_ap,Chl,green_= *E*_ap,Chl_ / *f*_green_. Other area-based inputs such as maximum carboxylation rate *V*_cmax_ [μmol m^-2^ s^-1^], as well as model outputs (e.g., internal CO_2_ concentration, *C*_i_ [μmol m^-3^]) refer to green leaves only. Assimilation (*A*_c_) is therefore initially computed per unit green leaf area. The stomatal conductance (*r*_cw_) as output of the leaf biochemical model is further used to calculate the transpiration of green leaves λ*E*_green_, also per unit green leaf area. Consequently, both fluxes first calculated per unit green leaf area, and later scaled with *f*green.

### 2.4 Fluorescence

SCOPE computes leaf level fluorescence emission using three main elements: incident irradiance in the excitation range 400-750 nm, excitation-fluorescence (E-F) matrices (*M*(λ_e_,λ_f_) and *M’*(λ_e_,λ_f_) for backwards and forward fluorescence, respectively), and the amplification factors *Φ’*_*f*_ which are provided by the biochemical model for sunlit and shaded leaves. E-F matrices represent the excitation of chlorophyll and the radiative transfer of incident and re-emitted radiation inside the leaf (Vilfan et al. 2016). In senSCOPE the leaf fluorescence emission is only calculated for green leaves, because for senescent leaves, the E-F matrices equal zero. Then the emission is scaled with *f*_green_.

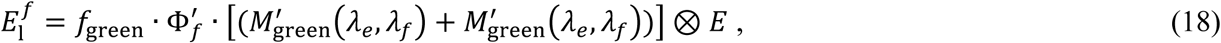

Leaf level fluorescence emission is then propagated to top of the canopy combining the same radiative transfer approach used by SCOPE and the averaged leaf optical properties (*ρ* and *τ*) for the mixed canopy.

### 2.5 Xanthophyll cycle

A recent version of SCOPE incorporates Fluspect-CX (Vilfan et al. 2018), a leaf RTM that simulates the variations in leaf optical properties induced by the activation of the xanthophyll cycle for photosynthetic down-regulation-, and propagates these variations from leaf to canopy level radiances. Changes in leaf optical properties are computed after photosynthesis, as a function of the rate coefficient for non-photochemical quenching (*K*_n_) provided by the biochemical module. This rate serves as a scaling factor of leaf *ρ* and *τ* between two extreme cases of with completely activated and completely deactivated xanthophyll cycle. In senSCOPE, senescent leaves show no carotenoids, no xanthophyll cycle and no related changes in optical properties; for this reason, the extreme cases calculated on the averaged *ρ* and *τ* simulate only variations induced by the green leaves. *K*_n_ is a rate defining the probability of the different fates of photons exciting chlorophyll, therefore, and similarly to *Φ’*_*f*_, it does not require additional correction. Therefore, senSCOPE uses the same radiative transfer functions than SCOPE for the propagation of signals related with the xanthophyll cycle.

## 3. Methods

### 3.1 Comparison with SCOPE model. Sensitivity analysis

In order to evaluate the differences between senSCOPE and the original model SCOPE (van der Tol et al. 2009), we run two different series of forward simulations modifying separately meteorological variables (F_meteo_) and vegetation properties (F_veg_). Eleven different canopies with *f*_green_ ranging between 0.0 and 1.0 with steps of 0.1 were simulated. As input to SCOPE we used weighted averages of the leaf parameters of each leaf type; similarly as field leaf measurements would be averaged to calculate canopy mean values.

In order to provide realistic meteorological forcing in the simulations F_meteo_, we used actual measurements acquired in the research site of Majadas de Tiétar between 5^th^ and the 20^th^ May 2016 (day of the year (DoY) 126 and 141, respectively). Fig. 2 summarizes this dataset.

**Figure 2.**
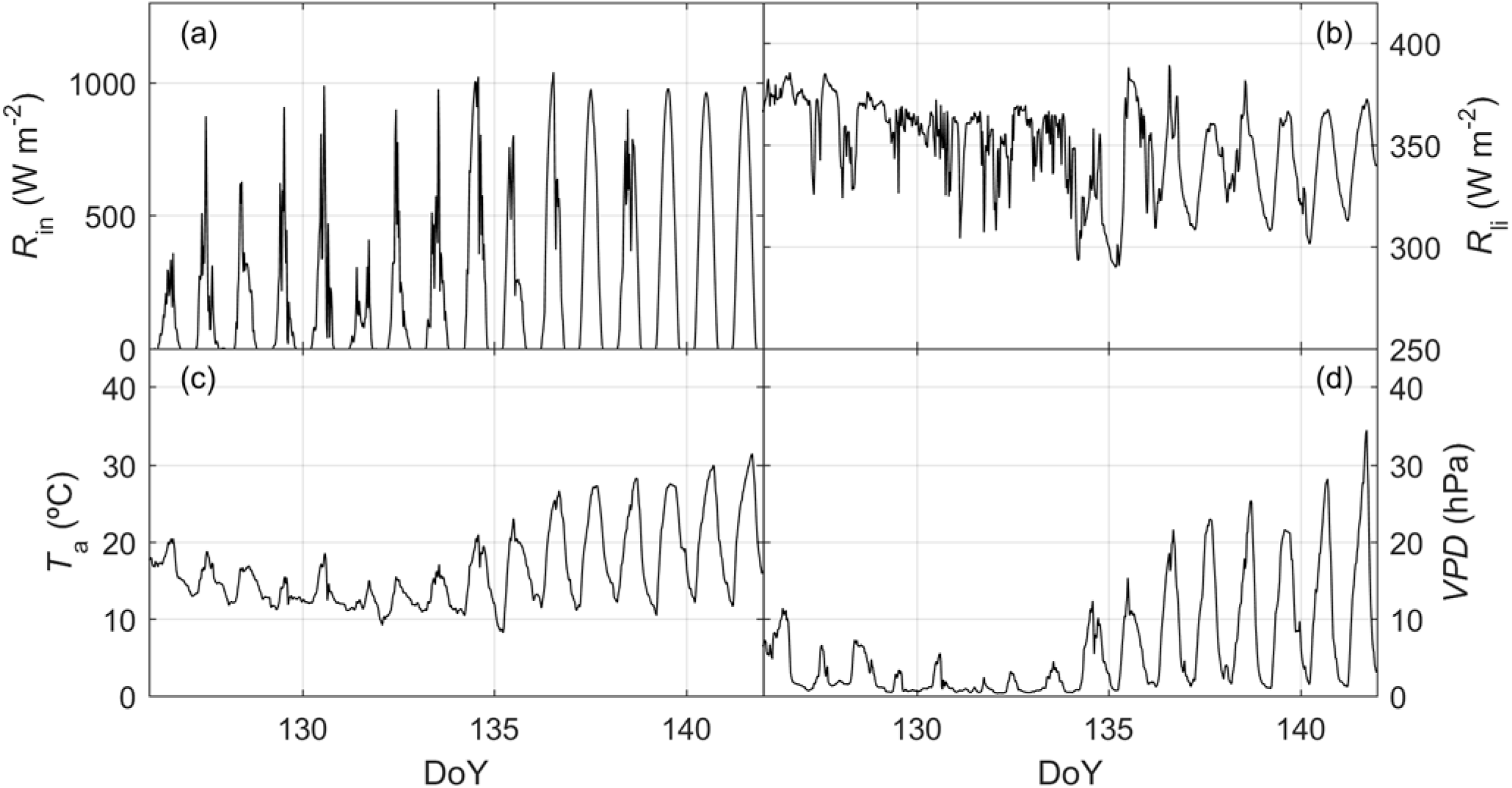
Short wave (a) and long wave incoming radiation (b), air temperature (c) and vapour pressure deficit (d) recorded in Majadas de Tiétar between the 5^th^ and the 20^th^ May 2019 (DoY 126 and 141, respectively) used in the forward simulation F_meteo_.

Short wave incoming radiation (*R*_in_, W m^-2^), long wave incoming radiation (*R*_li_, W m^-2^), air temperature (*T*_a_, °C), atmospheric vapour pressure (*e*_a_, hPa), wind speed (*u*, m s^-1^), air pressure (*p*, hPa) and soil moisture (*SM*_p_, % volume) were provided by a sub-canopy eddy covariance station at 1.6 m height (detailed description of the system can be found in El-Madany et al, (2018) and Perez-Priego et al, (2017)). Vapour pressure deficit (*VPD*, hPa) was calculated from *T*_a_ and *e*_a_; also, soil resistance for evaporation from the pore space (*r*_ss_, s m^-1^) was estimated as a function of using *SM*_p_ the model SCOPE v1.73. Sun zenith (*θ*_s_) and azimuth (*φ*_s_) angles were computed from timestamps and site location. In the F_meteo_ runs, only the abovementioned variables were modified; leaf and canopy properties were kept constant for the different *f*_green_ levels tested. Only daytime data (*θ*_s_ < 85.0 deg) were used in the simulation; which equals 422 runs per model and *f*_green_ level.

F_veg_ represented varying vegetation properties under constant meteorological conditions. To do so, we selected midday conditions of the 18^th^ May 2019 (DoY 139). A look up table with 500 samples of *C*_ab_, carotenoids concentration (*C*_ca_), *V*_cmax_, Fluorescence quantum efficiency (*f*_qe_), *m* and *LAI* was generated using Latin Hypercube Sampling (McKay et al. 1979). *C*_ca_ and *V*_cmax_ were constrained as a function of *C*_ab_ mimicking the relationships (linear function and noise) reported in Sims and Gamon (2002) and Croft et al, (2017), respectively. Table 1 shows the ranges of variation generated for each parameter varying in each F_veg_ simulation. Additionally, a smaller dataset was generated modifying only *LAI* or *C*_ab_ (and *V*_cmax_ and *C*_ca_ as a function of these) to illustrate an example of the response of models to these parameters. Several model outputs and internal parameters were evaluated. Moreover, we also compared the predicted underlying water use efficiency (*uWUE*, Eq. 19):

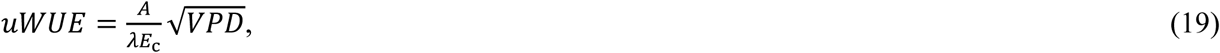

where *λE*_c_ is the canopy *λE*, excluding evaporation from the soil.

**Table 1.**
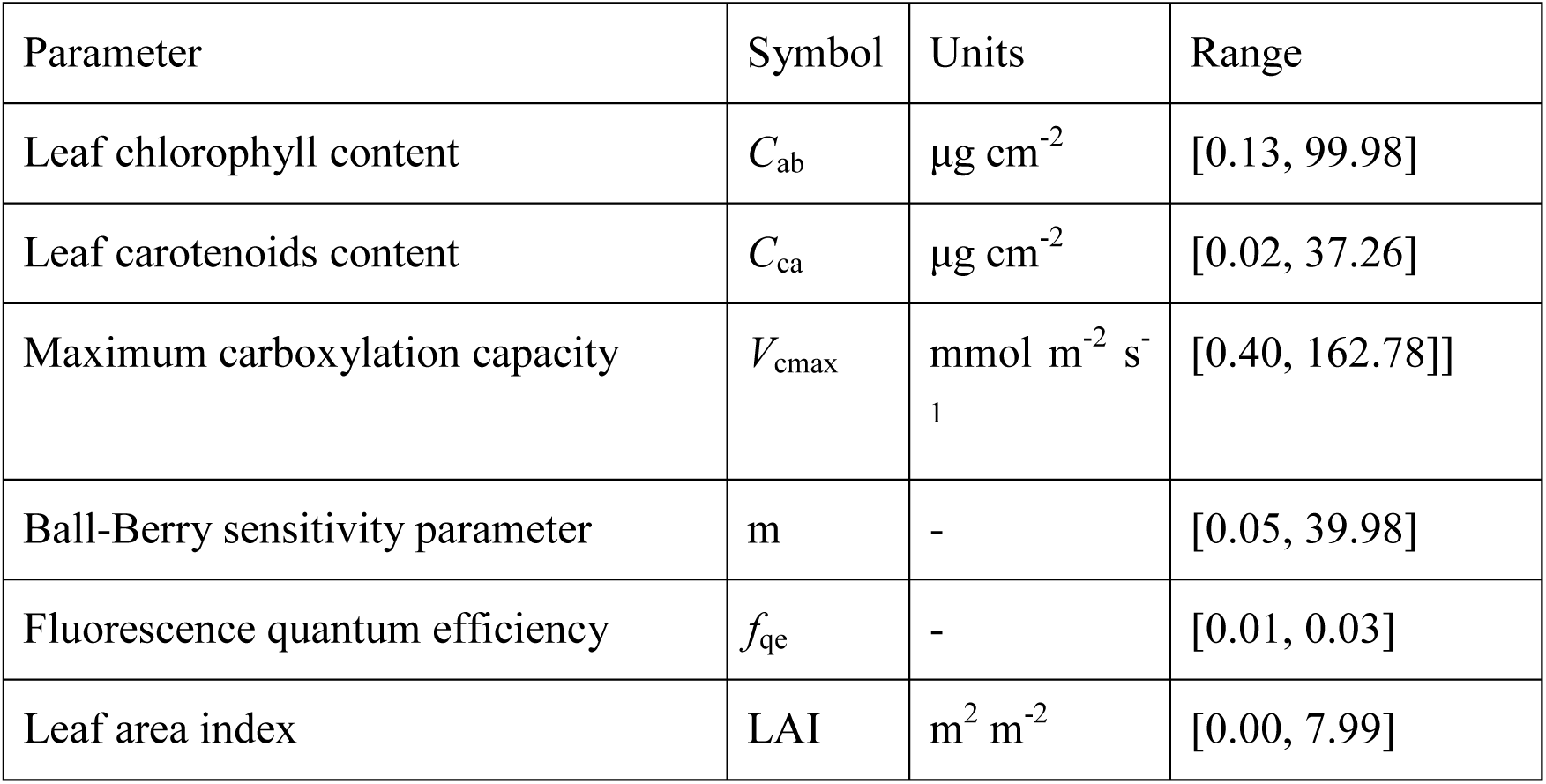
Vegetation parameters used in the forward simulation F_veg_.

| Parameter | Symbol | Units | Range |
| --- | --- | --- | --- |
| Leaf chlorophyll content | $C_{ab}$ | $\mu\text{g cm}^{-2}$ | [0.13, 99.98] |
| Leaf carotenoids content | $C_{ca}$ | $\mu\text{g cm}^{-2}$ | [0.02, 37.26] |
| Maximum carboxylation capacity | $V_{cmax}$ | $\text{mmol m}^{-2} \text{ s}^{-1}$ | [0.40, 162.78]] |
| Ball-Berry sensitivity parameter | $m$ | - | [0.05, 39.98] |
| Fluorescence quantum efficiency | $f_{qe}$ | - | [0.01, 0.03] |
| Leaf area index | LAI | $\text{m}^2 \text{ m}^{-2}$ | [0.00, 7.99] |

The Matlab^TM^ Profiler (Matwoks Inc., Natick, MA, USA) was used to evaluate the computing time and number of calls of the different functions of each model used during these simulations in each run. These metrics, together with the total computation time and the number of unsuccessful runs -where the energy balance does not succeed to converge to a solution-, were used to compare models’ performances.

### 3.2 Comparison with SCOPE model. Forward simulation with observational datasets

SCOPE and senSCOPE were also run forward using observational datasets from the study site of Majadas de Tiétar, Cáceres, Spain (39° 56’ 24.68″N, 5° 45’ 50.27″W). Observations -and when missing estimates-of vegetation properties and forcing variables integrated at ecosystem scale were used to run both models. Predicted fluxes and reflectance factors where compared with EC observations and hyperspectral airborne imagery.

#### 3.2.1 Study site and datasets

The study site is located in the experimental station of Majadas de Tiétar. It is a managed tree-grass ecosystem combining sparse trees (*Quercus ilex* L. subsp. *ballota* [Desf.] Samp) and a highly diverse herbaceous cover combining numerous species of three main functional plant forms: grasses, forbs and legumes. The climate is continental Mediterranean so that the grassland shows a strong seasonality initiated by greening phase around April, followed by a dry season that starts between May and June, a second re-greening driven by autumn rains, and a dormant phase during winter (El-Madany et al. 2018). The grassland phenology and functioning strongly responds to light and temperature in spring and to water availability in late spring-summer and in autumn (Luo et al. 2018). Several species grow and senesce at different times, usually, in early spring senescent material remnant from the winter is already present, then new material is also generated during spring, where *f*_green_ can already be already as low as ∼0.7 (Melendo-Vega et al. 2018).

In this site, three EC towers monitor three areas of the same ecosystem, one of them fertilized with nitrogen (N) and another one with N plus phosphorous (P), and the control one with no fertilization. These towers include also eddy covariance systems around 15 m above the ground, providing ecosystem-level measurements of carbon and water fluxes. Three sub-canopy towers monitor grassland fluxes ∼1.6 m aboveground. Details of the instrumentation and the manipulation can be found in El-Madany et al, (2018) and Perez-Priego et al, (2017). Also, a series of airborne campaigns with the Compact Airborne Spectrographic Imager CASI-1500i (Itres Research Ltd., Calgary, AB, Canada), operated by the Instituto Nacional de Técnica Aeroespacial (INTA) were conducted between 2012 and 2017. From a total of 17 images, a *R* of the footprint of each EC tower and campaign was extracted. Details of methods and data processing can be found in Pacheco-Labrador et al., (2017). Additionally, in each of the airborne campaigns, destructive sampling of vegetation provided estimates of ecosystem *LAI, f*_green_, *C*_dm_, *C*_w_, Nitrogen concentration (*N*_mass_) and/or *C*_ab_ and *C*_ca_. Further information on protocols and methods can be found in Melendo-Vega et al., (2018), Gonzalez-Cascon et al., (2017) and (Gonzalez-Cascon and Martin 2018).

#### 3.2.2 senSCOPE and SCOPE. Forward simulation and evaluation

Observed/estimated forcing variables and vegetation properties were used to predict fluxes and reflectance factors ±1 day around each flight campaign in each EC tower during daytime. Since no field observations of all the vegetation parameters were available, some of them had to be estimated. When missing, *C*_ab_ and *C*_ca_ were estimated from their relationship with *N*_mass_ observed in the site. Also *V*_cmax_ was estimated as a function of *N*_mass_ in the green leaves (*N*_mass,green_) following the relationship in Feng and Dietze (2013), and assumed 45 μmol m^-2^ s^-1^ for tree leaves. A constant *m* parameter of 10 was assumed, *N* and *LAD* were assumed 1.5 and spherical, respectively. *C*_s_ was estimated from the remaining leaf parameters inverting the statistical model described section 3.3.2 and in Appendix A. Soil reflectance was determined by *SM*_p_ and the parameters estimated by inversion of the BSM model (Verhoef et al. 2018) in Pacheco-Labrador et al (2019). Also, *r*_ss_ was estimated as function of *SM*_p_ using the model in Pacheco-Labrador et al (2019).

Then, we evaluated the capability of both models to predict *GPP, λE, R*_n_, *G*, and *H* comparing SCOPE and senSCOPE predictions with EC fluxes in the site. We also evaluated model performance and structure using predicted fluxes and computing quantities that describe energy partitioning, the evaporative fraction (Eq. 20)

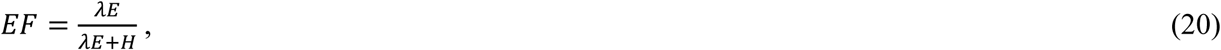

where *λE* and *H* are the total latent heat sensible heat fluxes, respectively.

Emitted irradiance in the TIR (*E*_t_) was compared with net radiometer measurements in the EC towers (CNR4, Kipp and Zonen, Delft, Netherlands); also *R* were compared with those of the imagery at the time of the overpass.

### 3.3 Comparison with SCOPE model. Inversion on observational datasets

In order to assess the impact of accounting for senescence material during the estimation of key biophysical (e.g., *LAI, C*_ab_) and functional (e.g., *V*_cmax_, *m*) vegetation parameters, we compared the parameter estimates and posterior predictions resulting from the inversion of both models against real observations in a Mediterranean grassland in the context of a nutrient manipulation experiment with N and P, featuring *f*_green_ between 0.05-1. In this work, we inverted SCOPE and senSCOPE using the inversion method and approaches proposed in Pacheco-Labrador et al. (2019).

#### 3.3.1 Study site and datasets

The inversion the models is tested using field observations from the understory grass layer of the site of Majadas de Tiétar, Cáceres, Spain, acquired in the context of the Small-scale MANIpulation Experiment (SMANIE) (Perez-Priego et al. 2015). This manipulation nutrient experiment was performed in an open area to minimize the effect of trees. The experimental design consisted of 4 blocks (4 replicates each) with N, P, both (NP) additions, and the control treatment (C, not fertilized). As a result of the fertilization, N, NP and P treatments induced changes in plant community, plant structure and function (Martini et al. 2019; Migliavacca et al. 2017; Perez-Priego et al. 2015). 9 field campaigns took place between 2014-2016 covering spring and early summer. In each block, midday measurements were carried out in two different collars with a dual spectroradiometric system providing hyperspectral *R* and SIF estimates in the O_2_-A (*F*_760_) and the O_2_-B (*F*_687_, not in all the campaigns). Diurnal time course of TIR up-welling radiance (*L*_t_) and *GPP* was determined using gas exchange chambers from sunrise to sunset. Fluxes were collected quasi-simultaneously in the same collars of the radiometric measurements at midday. The mismatch between the radiometric and chamber measurements was minimum. Moreover, destructive sampling near by the collars provided estimates of plant traits (*f*_green_, *LAI*, and nitrogen concentration *N*_mass_). Additional information about instrumentation, sampling methods and data processing can be found elsewhere (Martini et al. 2019; Migliavacca et al. 2017; Pacheco-Labrador et al. 2019; Perez-Priego et al. 2015).

#### 3.2.2 senSCOPE and SCOPE. Inversion and evaluation

We inverted senSCOPE and SCOPE using the same datasets and methodology described for the inversion of SCOPE in Pacheco-Labrador et al. (2019). Observations of *R* and *L*_t_, *F*_760_ and/or *GPP* were used to estimate *LAI, C*_ab_, *V*_cmax_, *m* and other biophysical parameters (Table 2) using an innovative methodology that combined biophysical and functional constraints in two different steps. Three different sets of constraints (inversion schemes) were tested, each combined in the first step of the inversion (Step#1), noon *R* with noon *GPP* (I_GPP_), noon *GPP* and *F*_760_ (I_GPP-SIF_), or nothing else (I_R_).

**Table 2.**
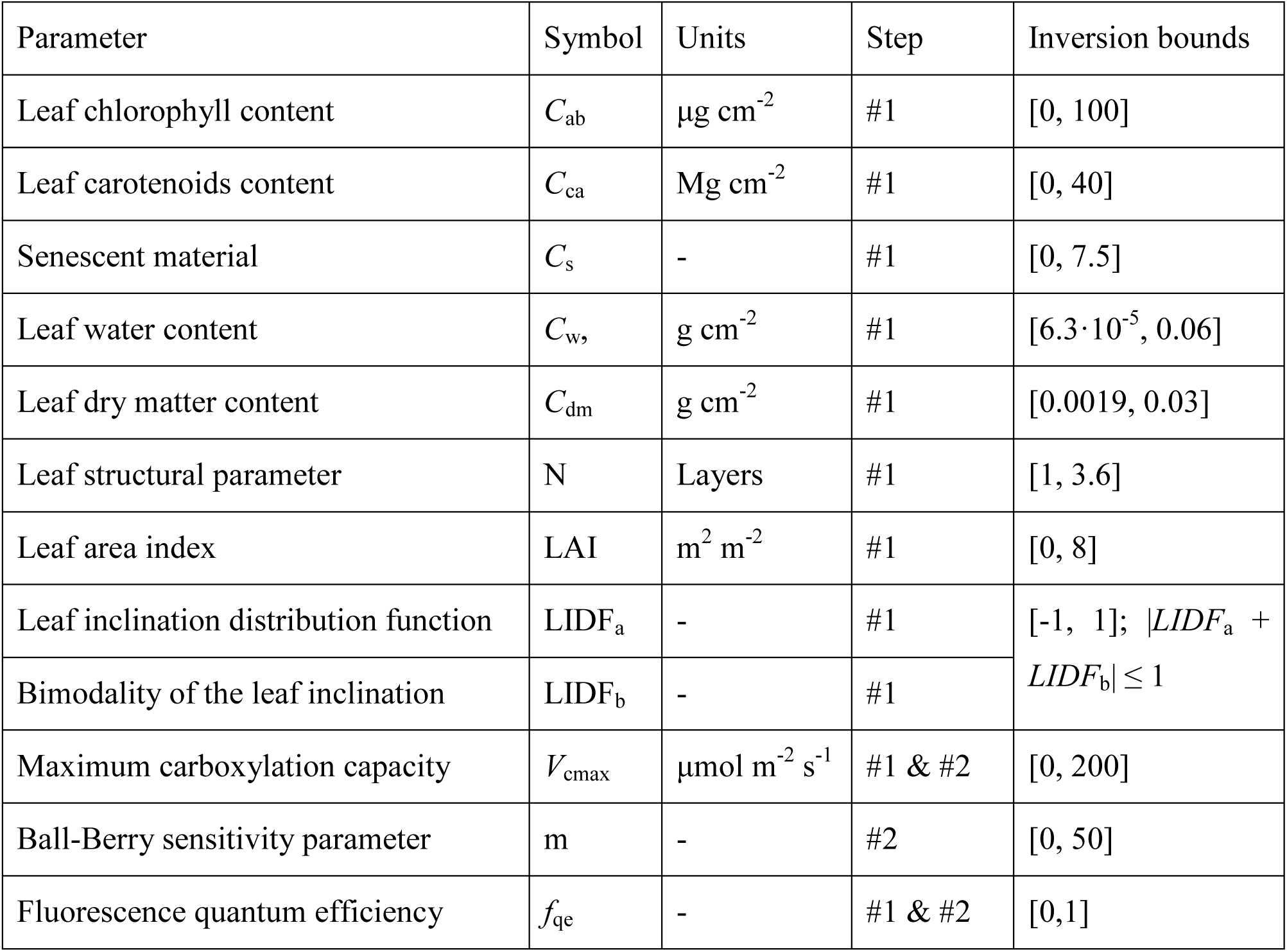
Parameters estimated inverting senSCOPE model.

In Step#1 biophysical parameters of the SCOPE model and a first guess of *V*_cmax_ were estimated. Uncertainties were estimated using a Bayesian approach (Omlin and Reichert 1999). Then, in a second step (Step#2) the guess of *V*_cmax_ was used as a prior and diel cycles of *L*_t_ combined with diel *GPP* (I_GPP_), diel *GPP* and noon *F*_760_ (I_GPP-SIF_), or only diel *L*_t_ (I_R_) were used to estimate the functional parameters *V*_cmax_ and *m. f*_qe_ was estimated in both steps in the schemes I_SIF_ and I_GPP-SIF_. Also, pattern-oriented model evaluation was used to assess the results of the different schemes. Unlike the previous work, this time we increased the inversion bounds (Table 2) for *C*_dm_ and *C*_w_ according to observed distributions in the site (Martín et al. 2019; Melendo-Vega et al. 2018). Also, since previous works found problems to cover the range of *R* in the near infrared, *C*_s_ upper bound was raised up to 7.5 a.u.; a value that allowed covering the low *R* values found in dry periods in the ecosystem (Martín et al. 2019). The multiple constraint inversion approach proposed in Pacheco-Labrador et al. (2019) provided coherent parameter estimates when *GPP* constrained the inversion (I_GPP_ and I_GPP-SIF_) using SCOPE; however, uncertainties in part related to the presence of senescent materials biased the estimation some of the parameters, notably *C*_ab_ during the dry season. In all the cases senescent material also was suspected to induce underestimation of *LAI*.

We used the same methodology to invert senSCOPE on the same datasets in order to compare the results provided by both models and to understand the suitability of using senSCOPE in environments featuring large fractions of senescent leaves. However, in the case of senSCOPE, 6 leaf parameters of two different leaf types must be estimated (Table 2). In order limit the number of free parameters in the inversion, we applied the following constraints: We assumed that green leaves presented no senescent pigments (*C*_s_ = 0) whereas senescent leaves only presented senescent pigments (*C*_ab_ = *C*_ca_ = 0). We also assumed that the mesophyll parameter (*N*) and dry matter content (*C*_dm_), were the same for both types of leaves, whereas that water content (*C*_w_) of green leaves was four times higher than senescent *C*_w_ (Kidnie et al. 2015). This allowed us reducing the degrees of freedom by 6. We assumed that average leaf parameters (*X*) could be computed as a linear combination of the parameters of each leaf type (*X*_green_ and *X*_senes_) as in Eq. 21:

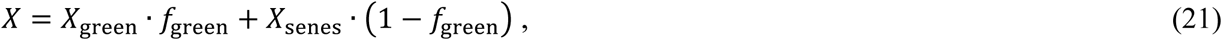

Given the constrains imposed on leaf parameters, we could directly optimize the leaf averaged parameters in the inversion, similarly as parameters are retrieved in the inversion of SCOPE (Pacheco-Labrador et al. 2019). To do so, *X*_green_ or *X*_senes_ are internally calculated solving them from Eq. 21; which is possible in all the cases since at least the value one of them together with *f*_green_ are known: Either they are equal, 0, or their ratio has been prescribed. senSCOPE includes the additional parameter *f*_green_; in order to reduce equifinality and as well as the number of parameters to estimate we prescribed *f*_green_ by modelling it as a function of the averaged leaf parameters *X* using a Neural Network (NN). The NN was trained from a look up table of individual *X*_green_ and *X*_senes_ parameters averaged as a function of *f*_green_; no assumptions on *N, C*_w_ and *C*_dm_ were made (Appendix A). As a result, the same parameters were estimated in the inversion of SCOPE and senSCOPE.

As in Pacheco-Labrador et al. (2019), we used pattern-oriented model evaluation approach to assess the retrieval of functional parameters, which cannot be determined from individual leaf measurements in the highly biodiverse grassland under study. To do so, we assessed the relationship of *V*_cmax_ and *C*_ab_ against *N*_mass_ in the green fraction of the canopy (*N*_mass,green_), and in the case of *V*_cmax_ it was compared with the relationship published by Feng and Dietze (2013) for grasslands. We also evaluated model performance and structure using not directly predicted fluxes, but variables derived from them such as *EF*, which describes energy partitioning (Eq. 19). In addition, a more traditional evaluation was also done assessing the goodness of the fit or prediction of model constraints (*R, L*_t_, *F*_760_, and/or *GPP*) and observed parameters (*LAI, f*_green_).

## 4. Results

### 4.1 Comparison of results and performance with SCOPE model: Sensitivity analysis

For the F_meteo_ runs, green and senescent leaf properties were kept constant for the different combinations of *f*_green_. Fig. 3a,b show the leaf optical properties simulated with senSCOPE and SCOPE, respectively. Accordingly Fig. 3c,d shows the TOC Hemispherical-Directional Reflectance Factors (*HDRF*) simulated with each model at midday of DoY 139, the timestamp used for F_veg_ runs. As can be seen, senSCOPE predicts spectroradiometric variables that vary more proportionally to *f*_green_, whereas SCOPE simulates stronger absorptions, especially in the visible region. This results from allocating all the absorptive substances to a single leaf type. The largest differences between models are found in the red and blue regions, where senescent leaves in senSCOPE increase scattering. We also verified that when *f*_green_ equals 1 or 0, the output of both models is the same.

**Figure 3.**
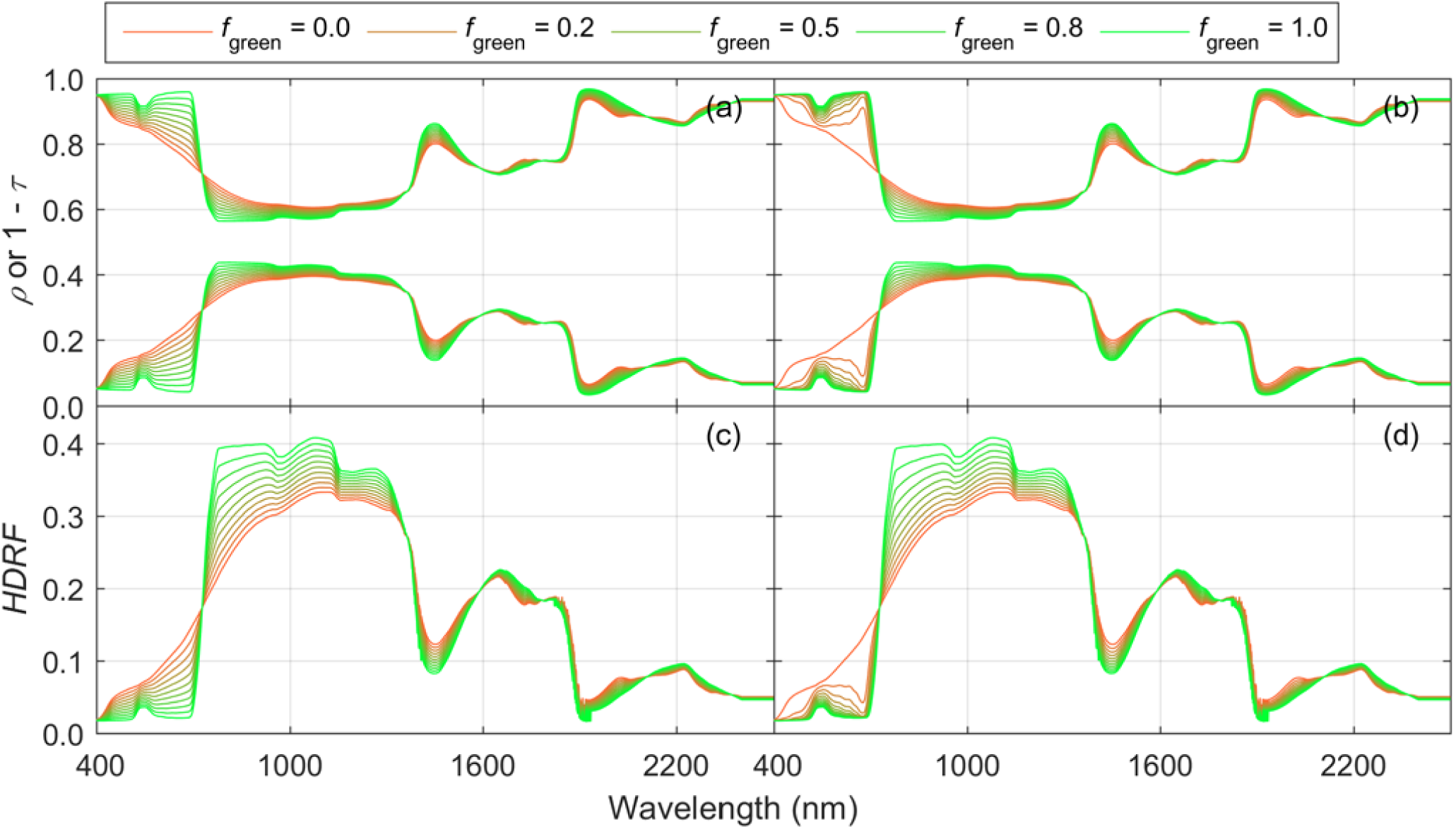
Leaf reflectance and transmittance factors predicted by senSCOPE (a) and SCOPE (b); and top of the canopy Hemispherical Directional Reflectance Factors predicted by senSCOPE (c) and SCOPE (d) for different fractions of green and senescent leaves.

Fig. 4 compares some of the spectoradiometric variables and fluxes predicted by senSCOPE (left semi-columns) and SCOPE (right semi-columns) during DoY 139 in the F_meteo_ runs. As can be seen, those variables that are strongly controlled by radiative transfer in the optical domain (*APAR*_Chl_ (Fig. 4c,d), the Photochemical Reflectance Index (*PRI*, Gamon et al, (1992)), sensitive to activation of the xanthophyll cycle (Fig. 4q,r) and *F*_760_ (Fig. 4s,t)) present a stronger and more linear sensitivity to *f*_green_. The same is observed for the water and energy fluxes (*λE* (Fig. 4e,f) and *H* (Fig. 4g,h)). Differences for variables related with the radiative transfer of thermal radiance seem to be lower (*R*_n_ (Fig. 4i,j) and *T*_c_ (Fig. 4k,l)). senSCOPE leaves feature a higher absorption of *PAR* per unit green leaf area, which produces a stronger NPQ activation (*K*_n_ (Fig. 4m,n)), and a depletion of photosynthetic efficiency around midday (*Φ’*_*f*_ (Fig. 4o,p)) for low *f*_green_ (unlike the other parameters, these are only representative of green leaves). Notice that the example shown is only representative of the meteorological and vegetation properties represented during DoY 139, and the differences shown should not be taken generally.

**Figure 4.**
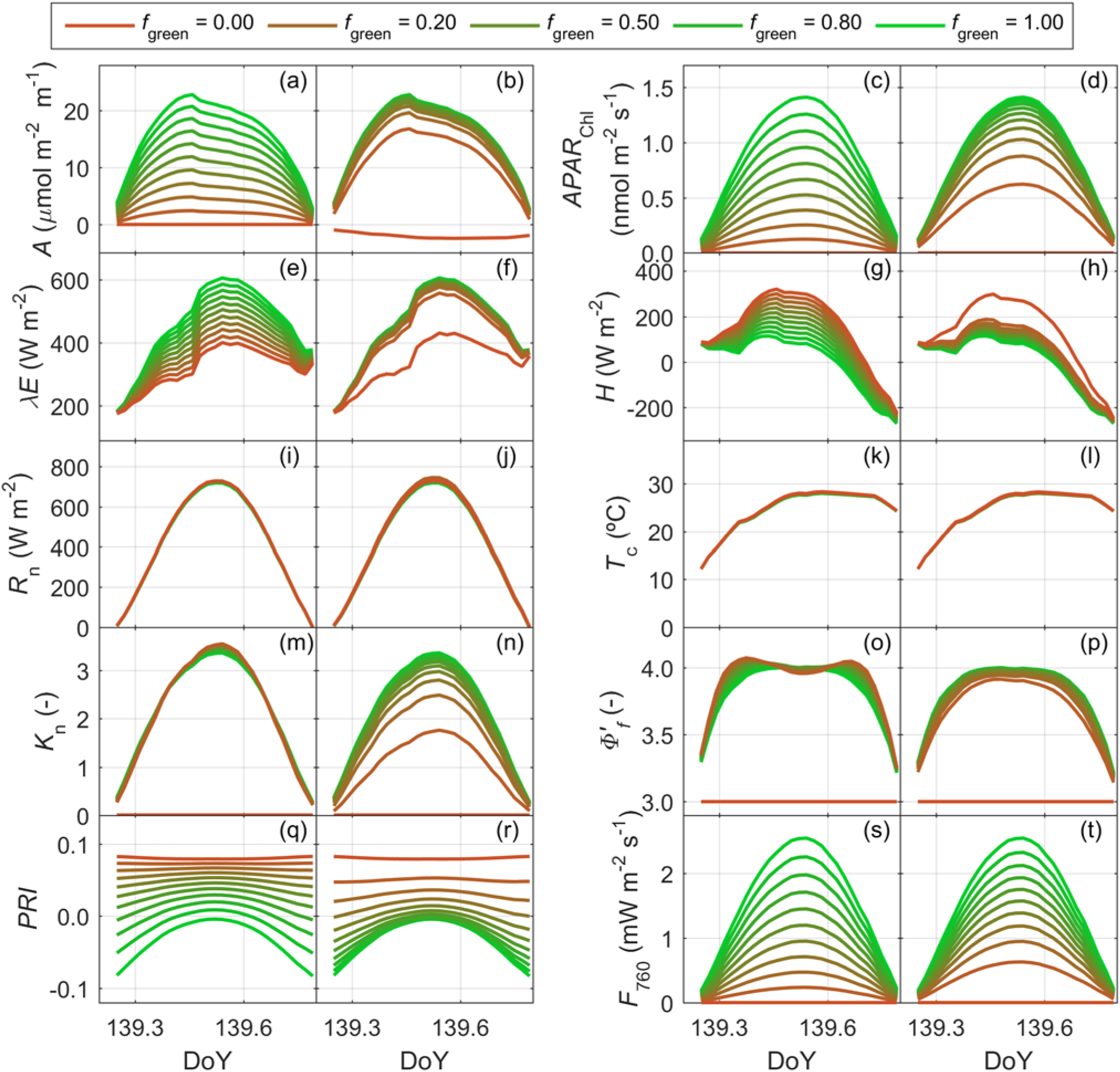
Diel cycles of senSCOPE (left semi-column) and SCOPE (right semi-column) predicted variables on DoY 136: Assimilation (a,b), photosynthetically active radiation absorbed by chlorophylls (c,d), latent (d,e) and sensible heat fluxes (g,h), net radiation (i,j), canopy temperature (k,l), rate coefficient for non-photochemical quenching (m,n), fluorescence efficiency (o,p), photochemical reflectance index (q,r) and TOC fluorescence radiance at 760 nm (s,t).

Fig. 5 shows the distributions of the difference between fluxes predicted by SCOPE and (minus) senSCOPE for each *f*_green_ level. Results of the F_meteo_ and the F_veg_ simulations are shown in the left and the right columns, respectively. As can be seen, both under varying meteorological conditions and varying plant properties, the two models predict the same fluxes when *f*_green_ = 1, but not always when *f*_green_ = 0. For *f*_green_ < 1 SCOPE predicts higher assimilation (*A*, Fig. 5a,b); but in the case of *f*_green_ = 0, where SCOPE predicts negative *A* due to photorespiration, and senSCOPE represents no photosynthetic leaf area. SCOPE also predicts in most of the cases higher *R*_n_ (Fig. 5c,d) and *λE* (Fig. 5e,f), and lower *H* (Fig. 5g,h) and *G* (Fig. 5i,j).

**Figure 5.**
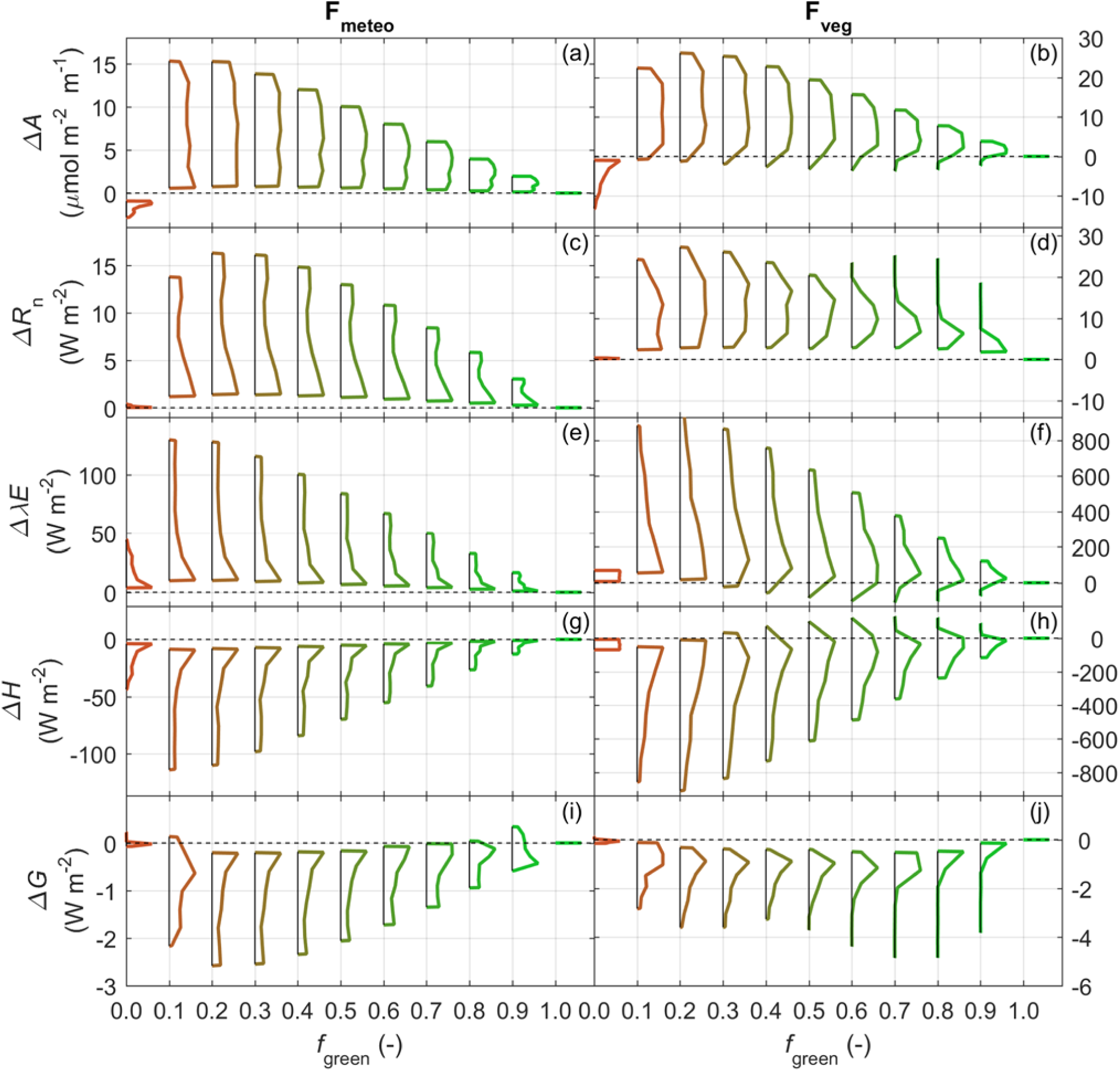
Distributions of the difference between the fluxes simulated with SCOPE and (minus) senSCOPE in the F_meteo_ run (left column) and the F_veg_ run (right column) for different fractions of green leaf area: assimilation (a,b), net radiation (c,d), latent heat flux (e,f), sensible heat flux (g,h) and soil heat flux (i,j).

Differences between predicted fluxes usually maximize when *C*_ab_ and *LAI* increase (Fig. S1a-e and S2 a-e, respectively). *G* (and *R*_n_) also present large differences for low values of these parameters and mid *f*_green_. In the analysis of the forward runs, the differences observed from F_veg_ simulations are often larger than those F_meteo_ simulations since the variability in the meteorological variables is -in relative terms-lower than the variability simulated for the vegetation properties.

For each *f*_green_ level, Fig. 6 presents the distribution of the difference between variables related to leaf function, as predicted by SCOPE and (minus) senSCOPE. Results of F_meteo_ and F_veg_ are shown on the left and the right columns, respectively. Similar to the fluxes, these variables are integrated according to *LAI* and the probability of each sunlit and shaded leaf angle. *APAR*_Chl_ (Fig. 6a,b) is equal for both models when the canopy is totally green or senescent. For the rest of the cases SCOPE predicts higher *APAR*_Chl_, except some cases when *C*_ab_ < 10 μg cm^-2^ (not shown). senSCOPE predicts higher canopy temperature (*T*_c_, Fig. 6c,d) than SCOPE; the largest differences are found when *C*_ab_ is high (Fig. S1g), or when *LAI* is low (Fig. S2g). Simlarly, *uWUE* (Fig. 6e,f) is higher for senSCOPE, but unlike *T*_c_ and most of the variables compared, differences in *uWUE* are more strongly controlled by meteorological conditions than by vegetation parameters. The largest differences in *uWUE* are found under cold conditions with *VPD* < 5 hPa (not shown). senSCOPE presents also higher *K*_n_ (Fig. 6g,h). Differences between models predictions increase with *LAI* (Fig. S2i), and decrease with *C*_ab_ (Fig. S1i). *Φ’*_*f*_ (Fig. 5i,j) is most often higher for senSCOPE than for SCOPE, especially if *LAI* is high and *C*_ab_ is low (not shown). On the other hand, SCOPE predicts higher *Φ’*_*f*_ when *LAI* is low (Fig. S2j) or when *C*_ab_ is low and *LAI* is moderate (Fig. S1j). As expected, both models predict the same values for these variables when *f*_green_ = 1.

**Figure 6.**
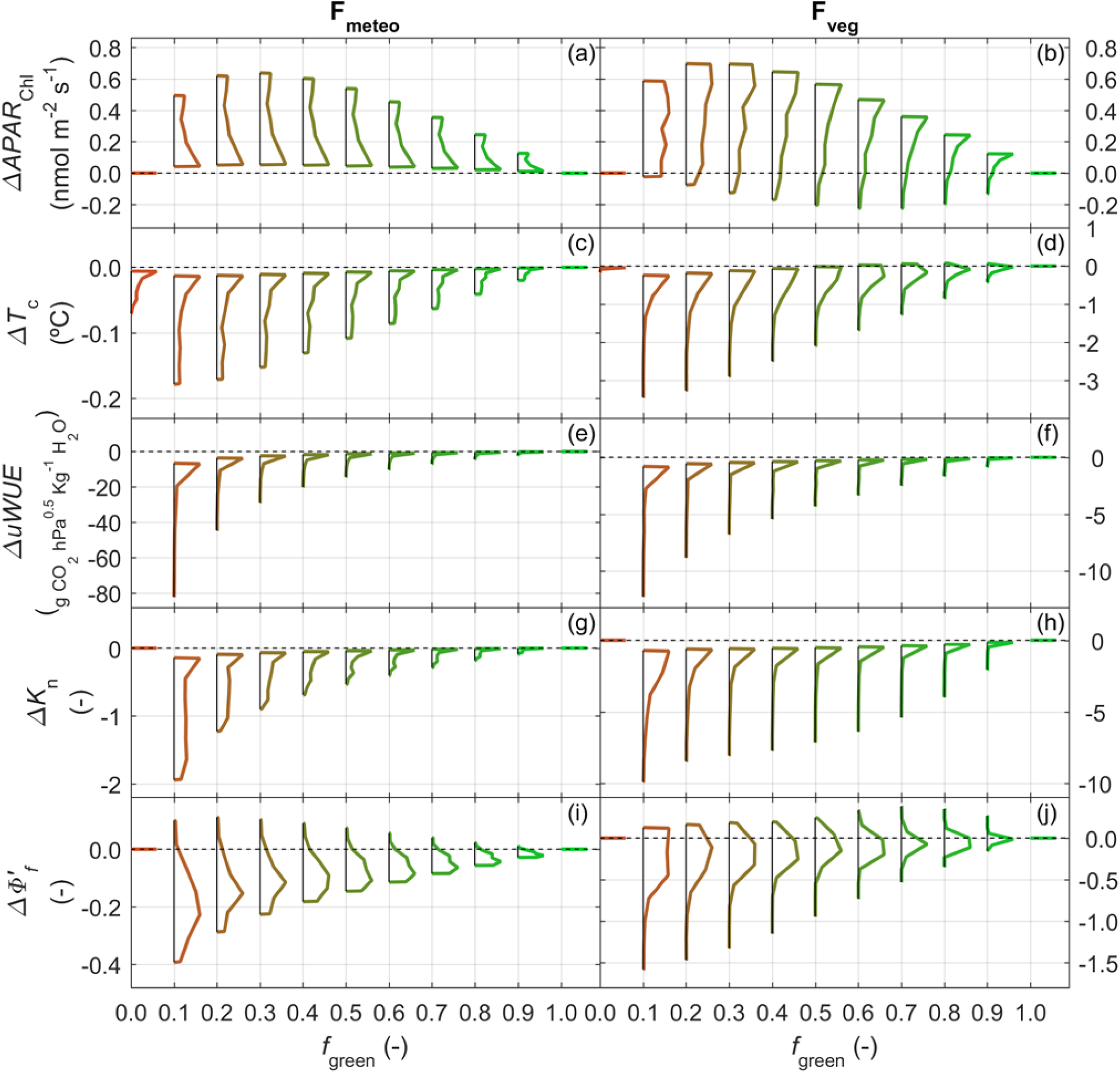
Distributions of the difference between variables indicative of plant physiology simulated with SCOPE and (minus) senSCOPE in the F_meteo_ run (left column) and the F_veg_ run (right column) for different fractions of green leaf area: photosynthetically active radiation absorbed by chlorophylls (a,b), canopy temperature (c,d), underlying water use efficiency (e,f), rate coefficient for non-photochemical quenching (g,h) and fluorescence efficiency (i,j).

Fig. 7 shows the distribution of some TOC spectroradiometric variables predicted by SCOPE and (minus) senSCOPE for each *f*_green_ level. Results of the F_meteo_ and the F_veg_ simulations are presented in the left and the right columns, respectively. *F*_687_ (Fig. 7a,b) and *F*_760_ (Fig. 7c,d) are larger for SCOPE in most of the cases the cases; the largest differences are found for low *f*_green_ and large *C*_ab_ (Fig. S1k,l) and LAI (Fig. S2k,l). Differences in *PRI* are negative for Fmeteo, but of both signs for Fveg (Fig. 7e,f). In this case, the influence of vegetation parameters is more complex and less linear than in other variables; since it depends on the combination of *C*_ab_ and *C*_ca_, their ratio and *LAI* (not shown). A similar analysis carried out on the *PRI* computed from reflectance factors where the effect of the xanthophyll cycle is not simulated reveals that differences between models rather respond to biophysical properties than to differences in function (not shown). Two more spectral indices responsive to pigments content and canopy structure are also analysed. Fig. 7g,h presents the differences for the Normalized Difference Vegetation Index (*NDVI*, Rouse et al, (1974)); Fig. 7i,j present differences for the MERIS Terrestrial Chlorophyll Index (*MTCI*, Dash and Curran (2007)); senSCOPE predicts lower and higher values, respectively. For these indices, the absolute difference between models increase as *f*_green_ decreases, and as *C*_ab_ and *LAI* increase (Fig. S1n,o and S2n,o, respectively).

**Figure 7.**
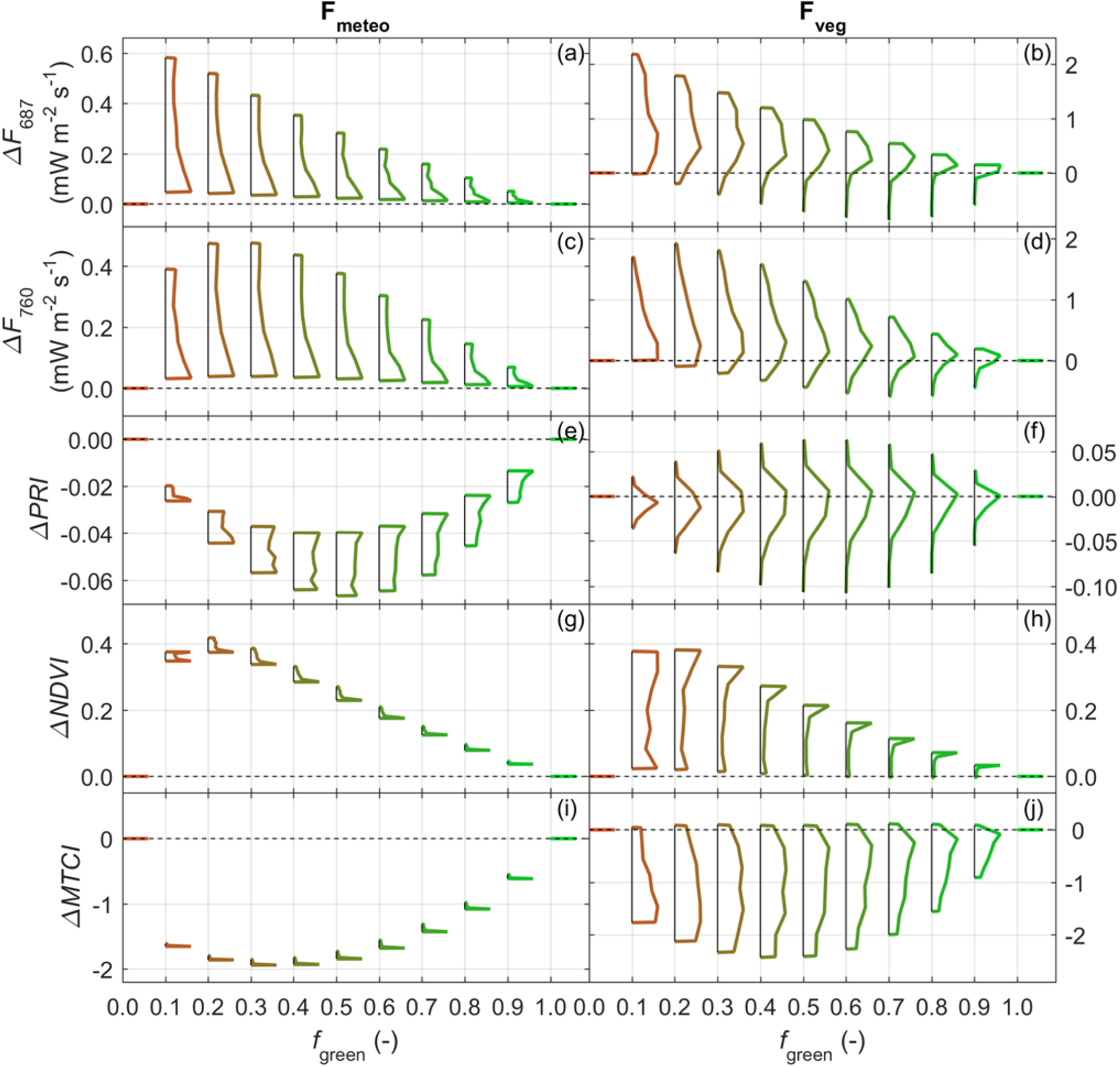
Distributions of the difference between spectral variables indicative of plant physiology, structure and biochemical composition simulated with SCOPE and senSCOPE in the F_meteo_ run (left column) and the F_veg_ run (right column) for different fractions of green leaf area: TOC fluorescence radiance at 687 nm (a,b), TOC fluorescence radiance at 760 nm (c,d), photochemical reflectance index including effects of xanthophyll cycle (e,f), normalized difference vegetation index (g,h) MERIS terrestrial chlorophyll index (i,j).

### 4.2 Comparison with SCOPE model. Forward simulation with observational datasets

Fig. 8 compares the different variables predicted by SCOPE and senSCOPE vs. the fluxes measured by the EC towers and *R* acquired by the airborne hyperspectral imager in the site of Majadas de Tiétar. The comparison is done using Total Least Squares (Golub and Loan 1980). In general, senSCOPE achieves higher coefficients of determination (*R*^2^) n lower relative root mean squared errors (*RRMSE*). Both models overestimate high *R* (Fig. 8a), and *GPP* (Fig. 8b); but senSCOPE is less deviated. SCOPE overestimates *λE* and *EF*, and underestimates *H* more than senSCOPE. Both models predict *R*_n_ quite accurately and precisely; but senSCOPE predicts *R*_n_, *E*_t_ and *G* with slightly larger errors and in some cases lower *R*^2^ than SCOPE.

**Figure 8.**
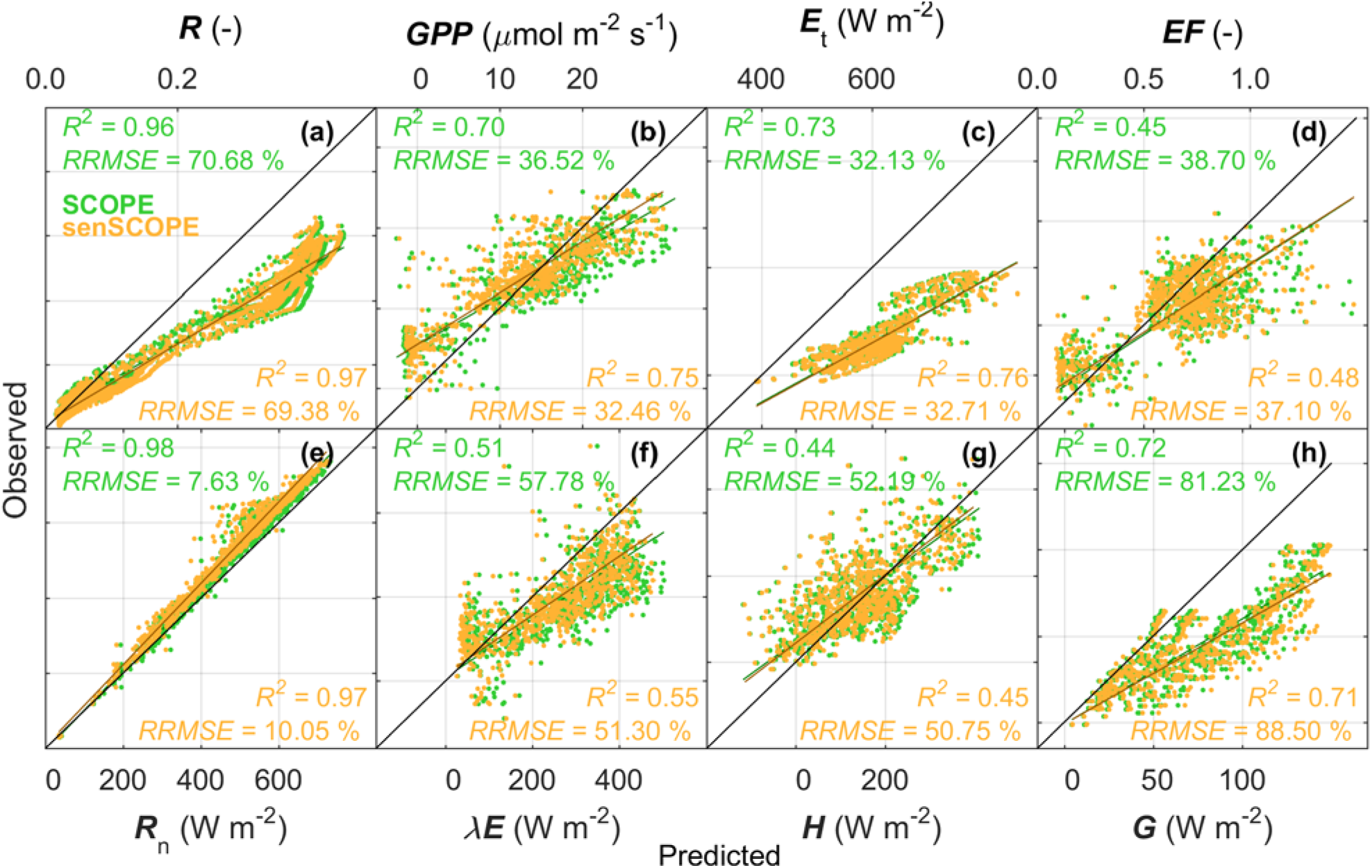
Comparison of observed and predicted fluxes and reflectance factors at ecosystem scale. Predictions are done by SCOPE (green) and senSCOPE (orange) using field observations or estimates of vegetation properties, as well as forcing variables measured at the research station of Majadas de Tiétar ±1 day around different airborne campaigns.

### 4.3 Comparison with SCOPE model. Inversion on observational datasets

Fig. 9 summarizes the capability of SCOPE and senSCOPE to fit/predict the variables used as inversion constraints in the different schemes tested; notice that not all the constraints are used to optimize parameters in all the schemes. The relative differences between the statistics of the fit are calculated as (100 · (*x*_senSCOPE_ - *x*_SCOPE_)/ *x*_SCOPE_); where *x* is the statistic and the respective model is presented in the subscript. *R*^2^ is estimated using Total Least Squares (Golub and Loan 1980), and the relative root mean squared error (*RRMSE*) and mean average error (MAE) result of the comparison of the observed/predicted values. Posterior uncertainty (*σ*_post_) is estimated according to Omlin and Reichter (1999). The relative differences of *R* in the visible spectral region (*R*_Vis_, Fig. 9a-d) and the near infrared (*R*_NIR_, Fig. 9e-h), *GPP* (Fig. 9i-l), *F*_760_ (Fig. 9m-p) and *L*_t_ (Fig. 9q-t) are presented for the three different inversion schemes tested. *R* is used in all the inversion schemes. senSCOPE fits *R*_Vis_ and *R*_NIR_ more poorly than SCOPE in the schemes I_GPP_ and I_GPP-SIF_; whereas in the case of I_R_ senSCOPE these are better fit and posterior uncertainties are lower than for SCOPE. senSCOPE slightly improves the fit of *GPP* when this is a constraint of the inversion; however, *σ*_post_ almost duplicate (values ∼80%, out of the plot scale). As in Pacheco-Labrador et al., (2019) I_R_ fails to accurately fit *GPP*, but *σ*_post_ is lower for senSCOPE. senSCOPE fit of *F*_760_ improves respect to SCOPE when this is a constraint of the inversion (I_GPP-SIF_), but *σ*_post_ increase in all the cases. senSCOPE fits *L*_t_ more poorly than SCOPE, but *σ*_post_ decrease in all the cases.

**Figure 9.**
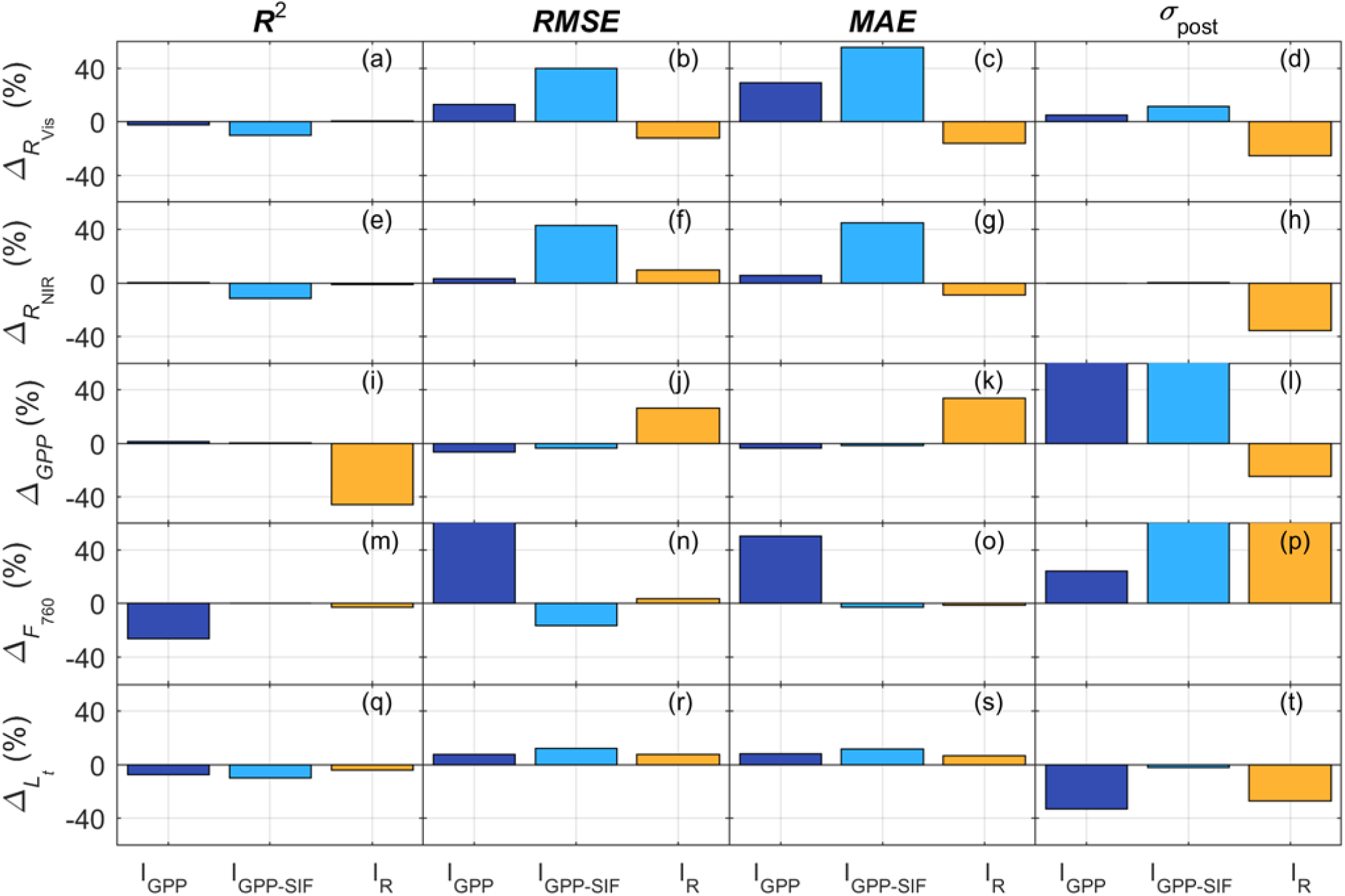
Relative difference between the fit/prediction statistics of the inversion constraints obtained by senSCOPE and SCOPE for the different inversion schemes.

Fig. 10 compares the most relevant model parameters estimated by SCOPE and senSCOPE for the different inversion schemes tested (presented by columns, from left to right: I_GPP_, I_GPP-SIF_, I_R_). Parameters are evaluated both using field observations and pattern-oriented model evaluation approach. *LAI* (Fig. 10a-c) and *f*_green_ (Fig. 10d-f) are compared against observations using Total Least Squares (Golub and Loan 1980). As can be seen, senSCOPE predicts similar *LAI* values but show higher *R*^2^ and significance. senSCOPE is also capable of providing reasonable estimates of *f*_green_, these are often overestimated but still within the bounds of the relationship *C*_ab_-*f*_green_ observed in the site (Fig. S3). *N*_mass,green_ is used to evaluate *V*_cmax_ (Fig. 10g-i) and to compare the relationship between both variables with the one reported in the literature for grasslands (Feng and Dietze 2013). Notice that senSCOPE *V*_cmax_ is provided per unit green leaf area and is thus comparable with SCOPE estimates and the literature data. Results are coherent with those presented in Pacheco-Labrador et al., (2019), I_R_ fails to constrain *V*_cmax_, whereas the schemes using *GPP* provide relationships with *N*_mass,green_ which are closer to those in the literature. *V*_cmax_ estimates are very similar for both models in I_GPP_; however, the use of *F*_760_ in I_GPP-SIF_ seems to slightly deviate the adjusted logarithmic model from the one fit to the data in Feng and Dietze (2013). Similarly, *C*_ab_ (per total leaf area) is evaluated against *N*_mass_ of the whole canopy (Fig. 10j-l), and their relationship is compared with field observations of both variables in the site of Majadas de Tiétar. When *GPP* constrains the inversion senSCOPE and SCOPE estimates are similar and follow the relationship observed in the field. However, as in Pacheco-Labrador et al., (2019), SCOPE I_GPP_ and I_GPP-SIF_ estimates present high values during the dry period, which stand out of the relationship with *N*_mass_ between 0.5-1.3 %. senSCOPE corrects most of these values, especially in the scheme I_GPP_; while the scheme I_GPP-SIF_ still preserves some of these high values. Fig. 10m-o compares predicted and observed *EF* using Total Least Squares (Golub and Loan 1980). In general, both models achieve similar results when *GPP* constrains the inversion; however, senSCOPE *R*^2^ are lower than in SCOPE. As in Pacheco-Labrador et al., (2019), I_R_ fails to constrain functional parameters.

**Figure 10.**
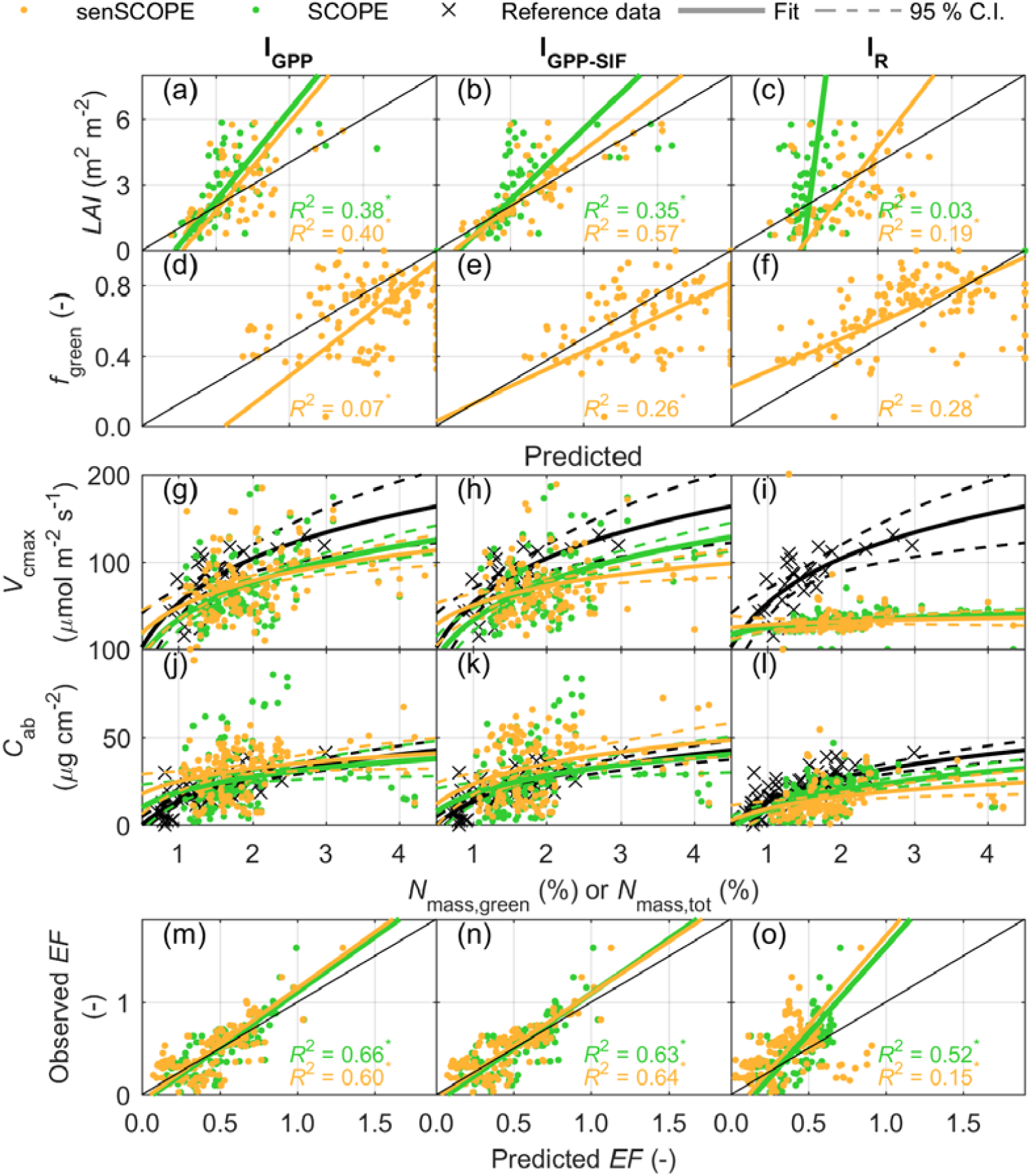
Summary of the parameters’ evaluation using observations and pattern-oriented model evaluation for the four inversion schemes tested. Leaf area index, (a-c) and green fraction of leaf surface (d-f), and evaporative fraction (q-t) are compared with field observation using Total Least Squares (Golub and Loan 1980). Significance is described with the symbols ^•^ for *p*-values 0.05≤ *p* <0.10; and ^*^ for *p* <0.05. The 1:1 line is shown in black. Maximum carboxylation rate (g-i) and chlorophyll concentration (j-p) are evaluated against nitrogen content in green leaves and total nitrogen content, respectively and compared with data from the literature (Feng and Dietze 2013) the first, and relationships observed in the field, the second A logarithmic relationship is fit in both cases, the 95 % confidence interval is show with dashed lines.

## 5. Discussion

This manuscript describes and evaluates senSCOPE, a version of the model SCOPE representing separately radiative transfer and physiological processes of green and senescent leaves; which is relevant in canopies featuring important senescent leaf area fractions. senSCOPE is evaluated against SCOPE 1) by direct comparison of forward synthetic simulations, 2) by comparison of simulated and observed ecosystem-scale fluxes and reflectance factors, and 3) by evaluation of parameter estimates and predicted variables via inversion of the models against a comprehensive dataset including hyperspectral optical *R*, as well as *GPP*, SIF and TIR radiance. These data were collected in a fertilization experiment with varying nitrogen and phosphorous additions and degrees of water stress. Results show that in senescent canopies senSCOPE improves the forward modelling of radiative transfer, photosynthesis and fluxes; and that in inversion -if suitably constrained-, it improves the estimation of *C*_ab_. At the same time, the performance of both models is comparable when green leaves dominate.

senSCOPE distributes senescent and remaining pigments in two conceptual leaves (green and senescent) and predicts separately their respective optical properties, which are later combined. This approach was already proposed by Bach et al., (2001) and used in later works (Bach and Verhoef 2003; Houborg et al. 2009; Houborg et al. 2015; Houborg and McCabe 2016; Verhoef and Bach 2003). This dual-leaf approach generates averaged “brighter” leaves since not all the absorbent species are located in the same leaf (Fig. 3). This has relevant consequences for the canopy-RTM, especially in those spectral regions where senescent and the rest of the pigments overlap, and therefore for *APAR*_Chl_. senSCOPE produces reflectance factors and *APAR*_Chl_ that close-to-linearly vary with *f*_green_; whereas in the case of SCOPE, these variables vary logarithmically with *f*_green_ since leaves absorptivity saturate due to the large presence of pigments. This saturation, combined with the fact that *R*_NIR_ was overestimated during senescence, led to unrealistically high *C*_ab_ estimates during the dry period when a strong functional constraint –*GPP*– was used (Pacheco-Labrador et al. 2019). Notice that only the constraint *GPP* provided robust estimates of functional parameters. In the present work, we repeated the inversion of SCOPE allowing higher *C*_s_ than in Pacheco-Labrador (2019) since this allowed predicting low *R*_NIR_ values observed in the site (Martín et al. 2019). This approach improved the fit of *R*_NIR_ for all inversion schemes during SCOPE inversion (not shown), but did not solve the overestimation of *C*_ab_ in the most strongly constrained schemes (I_GPP_ and I_GPP-SIF_, Fig. 10a,b). senSCOPE fitted less precisely the inversion constraints, and in some cases posterior uncertainties increased due to the strong control that *f*_green_ has on most of the model outputs (Fig. 9). For I_R_ senSCOPE improved the fit of *R*, but the opposite occurred when *aPAR* was constrained by *GPP*, suggesting that the model might not still represent accurately the observed grassland However, senSCOPE led to *C*_ab_ values more soundly related with *N*_mass_ than SCOPE during the dry season (schemes I_GPP_ and I_GPP-SIF,_ Fig. 10j,k).

The fact that senSCOPE limits photosynthesis and transpiration to the green fraction results in a close-to-linear relation between *f*_green_ on the one hand, and *A* and *λE* on the other hand (Fig. 4 and 5). SCOPE predicts higher assimilation and transpiration unless *f*_green_ is very low (∼0); in that case *A* is negative while *λE* is still high. Contrarily, *R*_n_ and *G* predictions are similar for both models; also, differences in *H* are lower than for *λE*, but still in senSCOPE *H* varies more linearly with *f*_green_ than in SCOPE (notice that *f*_green_ is not a SCOPE parameter, but is used to average leaf parameters). In the forward simulation at ecosystem scale senSCOPE predicted most of the ecosystem fluxes better than SCOPE (Fig. 8). In this case we assumed a fixed value for *m*, which might be not completely realistic; however additional works at ecosystem scale have shown that senSCOPE can more robustly represent water use efficiency than SCOPE (not shown). In the inversion at plot level, senSCOPE predicted *GPP* better than SCOPE when used as constraint (Fig. 9k-o). In contrast, *EF* was predicted more poorly in all the schemes. senSCOPE assumes no transpiration from senescent leaves; however evaporation from their surface might be relevant when these are moisturized by dew or rainfall. Neither SCOPE nor senSCOPE represent that process and their use after such situations might result uncertain.

In inversion, both SCOPE and senSCOPE underestimated *LAI*, while senSCOPE overestimated *f*_green_ (Fig. 10a-h). As discussed in Pacheco-Labrador et al, (2019) and Melendo-Vega (2018), the optical properties of dry standing material might not be accurately described by RTM, leading to an overestimation of *R*_NIR,_ which seems to be counter-weighted in inversion by reducing *LAI*. In fact, inversion schemes using *GPP* (I_GGP_ and I_GPP-SIF_) improved the estimation of *LAI* since *GPP* demands higher *APAR*_Chl_ in exchange for increasing the fitting error of *R*_NIR_ (Pacheco-Labrador et al., (2019), this work). In senSCOPE, underestimation of *LAI* was also compensated also by overestimating *f*_green_. These facts suggest that the optical properties of the senescent material and/or the death standing material of this grassland (and likely other ecosystems) are not accurately represented, leading to biased estimates of some of the parameters. In fact, it was necessary increasing the upper bound of *C*_s_ to be able to predict low *R*_NIR_ in the dry season. We allowed *C*_s_ up to 7.5; whereas values up to 5.0 are reported in literature (Houborg and Anderson 2009). Too high *C*_s_ might have led to unrealistic representation of *ρ* and *τ* of senescent leaves, very dark in the visible region but also with low *R*_NIR_. In some cases SCOPE estimated *C*_s_ = 7.5, whereas senSCOPE predicted *C*_s_ < 5 in most of the cases (Fig. S4c). Apart from *LAI, C*_dm_ and *C*_w_, -which are weakly constrained because the spectroradiometric measurements did not include the short wave infrared range (SWIR)-, might have been affected by this problem. SCOPE and senSCOPE estimates of *C*_dm_ often hit the upper bound stablished from observations in the field. High *C*_dm_ also serves to reduce *R*_NIR_. In contrast, senSCOPE *C*_w_ estimates are less often saturated; *C*_w_ has little effect below 970 nm, but influences leaf optical properties in the SWIR. The relationship between *N, C*_dm_ and *C*_w_ of green and senescent leaves assumed during inversion might have contributed to increase the uncertainty of the parameter estimates; for example, it has been observed that leaf thickness decreases during senescence (Castro and Sanchez-Azofeifa 2008); whereas other works assign high N values to senescent leaves (Houborg et al. 2009). However, a balance between model error and equifinality must be also observed. Site-specific relationships between the parameters of each leaf type or relationships found in global databases could be used in the future to improve the representation of semi-arid canopies. senSCOPE does not include improved calibrated absorption coefficients or refractive indices to more realistically represent senescent leaves and death standing material, but it offers a formally more correct representation of mixed canopies. The model improves the representation of these canopies, which could be used in the future to calibrate or validate specific absorption spectra of senescent material. senSCOPE can also be applied to other canopies, such as crops and forests, which are characterized a senescent stage. Moreover, the approach adopted in senSCOPE could be similarly used to represent other mixed canopies combining plants with different biophysical properties and function, such as C3 and C4 species. An additional problem for the representation of mixed canopies would be the vertical distribution of the senescent material. The impact on the observed *R* and fluxes is unclear, and further research is needed in this direction. In such studies, senSCOPE could also be extended to other versions of SCOPE, such as mSCOPE (Yang et al. 2017) to describe the vertical distribution of senescent matter.

*f*_green_ is a critical parameter in senSCOPE, it strongly controls RTM and fluxes and increases equifinality of the inverse problem. Thus, the use of prior information about is this variable is strongly recommended during inversion. For this reason, in this work *f*_green_ was indirectly predicted from leaf parameter estimates using a NN while the model was inverted. The design of this model was critical to achieve acceptable results, and during training *C*_ab_ (and *C*_ca_) had to be limited to the ranges observed in the study site (up to ∼40 μg cm^-2^). During inversion higher *C*_ab_ values were allowed, but still, *C*_ab_-*f*_green_ estimates stood within or very close to the bounds observed and used to train the NN (Fig. S3)

As a result of the combination of changes in RTM and photosynthesis, not only carbon and water fluxes, but also photosynthetic efficiency and downregulation resulted modified (Fig. 6). On one side, senSCOPE tends to predict higher canopy temperatures than SCOPE, especially when *f*_green_ decreases. Senescent leaves are warmer than green leaves, but senSCOPE green leaves are not necessarily cooler than SCOPE leaves (not shown). Leaf temperature strongly influences photosynthetic efficiency and together with *APAR*_Chl_ on photosynthesis down-regulation. Fig. 4m,h show how senSCOPE diel cycles of *K*_n_ reach higher midday values than SCOPE. SCOPE predicts larger variability of *K*_n_ as a function of *f*_green_ under conditions of low illumination, whereas senSCOPE *K*_n_ varies more strongly with *f*_green_ under high temperature and irradiance conditions (not shown). Non-photochemical quenching has also different effects on the predicted *Φ’*_*f*_. For example, Fig. 4o,p show how senSCOPE predicts a decrease of this efficiency at midday whereas this is hardly noticeable for SCOPE. *K*_n_ and *Φ’*_*f*_ are fundamental variables to mechanistically interpret SIF signals to determine functional status of vegetation and stress (Frankenberg and Berry 2018; Porcar-Castell et al. 2014). Thus considering the differences shown, both models can lead to very different interpretations. Adequate representation of physiological processes and their drivers is fundamental to mechanistically interpret these signals; but also the representation of the spectral variables used to obtain information about these processes, such as fluorescence radiance or *PRI*. Similarly as *R*, spectral indices vary more linearly with *f*_green_ in senSCOPE than in SCOPE (e.g., Fig. 4q,r). Unlike other spectroradiometric variables, *PRI* show no clear differences between models (e.g. distributions of the difference centre around 0). PRI is known to result sensitive to pigments pool, ratio and to *LAI* (Gamon and Berry 2012; Garbulsky et al. 2011); results of this work also show that this index is also strongly sensitive to the presence of senescent material. The magnitude of SIF emissions is also modified by senSCOPE, which tends to predict less SIF when *f*_green_ decreases, (Fig. 7a-d).

In this study we compare the inversion of SCOPE and senSCOPE using the data and approaches of in Pacheco-Labrador et al, (2019), but allowing for higher values for *C*_s_ (as well as *C*_dm_ and *C*_w_). The wider parameter bounds did not change significantly the results obtained with SCOPE, and differences were mainly related to the use of senSCOPE; which improved the estimation of *C*_ab_ in the dry season. As with SCOPE, SIF (not shown) and *R* failed to constrain functional parameters (e.g., *V*_cmax_) and *LAI*; and only inversion schemes relying on *GPP* provided robust estimates. However with senSCOPE, the schemes relying on SIF reduced their performance respect to SCOPE. I_GPP-SIF_ fitted the inversion constraints more poorly, and could not correct high *C*_ab_ estimates during senescence as much as I_GPP_. This might be result from the use of large *C*_s_, which suggests further work is needed to more accurately characterize the optical properties of death standing and senescent material. Also for senSCOPE, functional parameters resulted insensitive to I_R_ constraints (partly due to inversion method, see Pacheco-Labrador et al, (2019)). Bayat et al., (2018) inverted SCOPE using *R* and found troubles to predict low *GPP* and λE in a grassland during senescence, which was corrected constraining the model with *R* and TIR radiance to reduce *V*_cmax_ during this period. Fig. 10i-l compares *V*_cmax_ estimates of both models; for senSCOPE *V*_cmax_ is presented respect to green leaf area, whereas in SCOPE, it is presented respect to total leaf area (all considered “green”). As can be seen, when adequately constrained estimates of both models are comparable. In senSCOPE *GPP* scales with *f*_green_, and *V*_cmax_ (in the green leaves) does not need to decrease to predict low assimilation.

senSCOPE is computationally more demanding (around 10% slower) than SCOPE since more processes and calculations are needed, and more iterations are required to close the energy balance (Table S5). However, senSCOPE seems more robust and provides lower energy balance closure error. Since performance of both models is similar for large *f*_green_, both models can be alternately used through the season according the presence of senescent material.

## 6. Conclusions

The combination of advanced radiative transfer models with models representing exchanges of matter and energy between vegetation, soil and atmosphere is bringing new opportunities to improve our understanding of ecosystem function from remote observations. For example, the model SCOPE is being used in the last years with this purpose. However, the accuracy with which these models represent reality limits their application; and ecosystem-specific features can bias results and their interpretation. In this context, we present the model senSCOPE; which adapts SCOPE radiative transfer, energy balance, photosynthesis and transpiration in homogeneous canopies with mixed green and dry leaves. The separated representation of green and senescent leaves significantly modifies the simulation of fluxes and spectra signals respect to a model featuring a single leaf with “averaged” properties. senSCOPE reflectance factors, carbon assimilation and water and energy fluxes linearly scale with *f*_green_; it also improves the prediction of these variables in forward simulations as well as the estimation of vegetation parameters, notably *C*_ab_, during the dry season. This is significant for the remote sensing of vegetation function of semi-arid ecosystems, and potentially for phenology monitoring. Despite the improvements, results suggest that not only model structure needs to be corrected; a more accurate characterization of the optical properties of senescent material in grasslands is still needed. The use of SCOPE and derived models is growing in the remote sensing community; however, further assessment of their performance to inform about plant function should be tested in different ecosystems. For example, the role of vertical and horizontal heterogeneity is still unclear. Robust evaluation, e.g. pattern-oriented model evaluation approach, would contribute to identify caveats and ecosystem-specific features that prevent accurate monitoring of their function; and that therefore, should be also represented.

## Appendix A: Green fraction Neural Network predictor

In senSCOPE inversion, the fraction of green leaf area in the canopy (*f*_green_) is estimated as a function of the canopy averaged leaf RTM parameters using a NN model trained form simulated data. Latin Hypercube Sampling was used to generate a look-up table (LUT) with 5000 samples of different leaf constituents (*C*_ab_, *C*_ant_, *C*_dm_, *C*_w_, *C*_s_), *N* and *f*_green_. *C*_ca_ was included in the LUT as a function of *C*_ab_ according to the relationship reported in Sims and Gamon (2002), and an uncertainty estimated in the relationship of ∼4.5 μg cm^-2^ according to field measurements was used to add Gaussian noise. The same bounds that were applied in inversion (Section 3.3.2) were used to design the LUTs; however *C*_ab_ and *C*_ca_ of green leaves were limited to 40 and 10 μg cm^-2^, respectively; according to field observations. LUT values were assumed to belong to pure green and senescent leaves, and averaged leaf parameters were mixed according with Eq. 21, assuming that in green leaves *C*_s_ = 0, and that in senescent leaves *C*_ab_ = 0, *C*_ca_ = 0, *C*_ant_ = 0. No additional assumptions about the values of the parameters of each leaf type and therefore the *N, C*_dm_, *C*_w_ were taken directly from the LUT.

A NN was trained using SimpleR (Camps-Valls et al. 2012) to predict *f*_green_ as function of the canopy averaged leaf RTM parameters. During the training, 60 % of the dataset was used for fitting and 40 % for testing. Performance statistics are presented in Table A1.

**Table A1.** Statistics of the fraction green leaf area (*f*_green_) Neural Network (NN) model.

| Dataset | $R^2$ | RMSE | ME | MAE |
| --- | --- | --- | --- | --- |
| Training | 0.818 | 0.123 | 0.005 | 0.093 |
| Validation | 0.718 | 0.230 | 0.158 | 0.180 |

## Code availability

senSCOPE code and further developments, as well as the code for the multiple constraint inversion of the model are publicly available at https://github.com/JavierPachecoLabrador/senSCOPE.

## Author contributions

JPL, TSEM, MM and CvdT designed the model. JPL and MM designed model evaluation. TSEM, AC, OPP, JG, PM, RGC, GM, MR and MM provided measurements of fluxes, plant parameters and spectral variables. JPL, CvdT, MM, OPP, JG, PM and RGC wrote the paper.

## Acknowledgements

JPL, MM and MR acknowledge the EnMAP project MoReDEHESHyReS “Modelling Responses of Dehesas with Hyperspectral Remote Sensing” (Contract No. 50EE1621, German Aerospace Center (DLR) and the German Federal Ministry of Economic Affairs and Energy). Authors acknowledge the Alexander von Humboldt Foundation for supporting this research with the Max-Planck Prize to Markus Reichstein; the project SynerTGE “Landsat-8+Sent inel-2: exploring sensor synergies for monitoring and modelling key vegetation biophysical variables in tree-grass ecosystems” (CGL2015-69095-R, MINECO/FEDER,UE); and the project FLUχPEC “Monitoring changes in water and carbon fluxes from remote and proximal sensing in Mediterranean ‘dehesa’ ecosystem” (CGL2012-34383, Spanish Ministry of Economy and Competitiveness). Authors are very thankful to the MPI-BGC Freiland Group and especially Olaf Kolle, Martin Hertel as well as Ramón López-Jiménez (CEAM) for technical assistance. We are grateful to all the colleagues from MPI-BGC, University of Extremadura, University of Milano-Bicocca, SpecLab-CSIC, INIA and CEAM which have collaborated in any of the field and laboratory works. We acknowledge the Majadas de Tiétar city council for its support.

